# Generalizing deep variant callers via domain adaptation and semi-supervised learning

**DOI:** 10.1101/2023.08.12.549820

**Authors:** Youngmok Jung, Jinwoo Park, Hwijoon Lim, Jeong Seok Lee, Young Seok Ju, Dongsu Han

## Abstract

Deep learning-based variant callers (DVCs) offer state-of-the-art performance in small variant detection from DNA sequencing data. However, their reliance on supervised learning and the subsequent need for extensive labeled data pose a potential hurdle to their generalizability across diverse sequencing methods with varying error profiles. Indeed, even minor discrepancies in error profiles can compromise the robustness of DVCs and impair the variant calling accuracy in the target sequencing method. To mitigate these challenges, we propose RUN-DVC, the first semi-supervised training approach for DVCs that presents two complementary training techniques to the conventional supervised training approach. RUN-DVC leverages semi-supervised learning techniques to learn error profiles from unlabeled datasets of the target sequencing method as well as a domain adaptation technique to aid semi-supervised learning by reducing the domain discrepancy due to different error profiles. We analyze and contrast RUN-DVC against the supervised training approach under various generalization scenarios using nine sequencing methods from Illumina, BGI, PacBio, and Oxford Nanopore sequencing platforms. Remarkably, RUN-DVC significantly improves the variant calling accuracy of DVC in the target sequencing method even with purely unlabeled datasets in the target domain and enables label-efficient generalization when partially labeled datasets are available. Our results suggest RUN-DVC is a promising semi-supervised training method for DVCs with the potential to broaden the use of DVC across diverse sequencing methods.

## Introduction

Variant calling in DNA sequencing data is a critical piece in medical genetics as well as in population genetics and functional genomics. These applications crucially rely on accurate variant calling, which is finding small variants such as single nucleotide polymorphism (SNP) and short insertion-deletion (INDEL) from sequencing data [1–3]. Both variant calling algorithms and sequencing technologies are yet imperfect and continually evolving [4–7]. Thus, developing variant callers in line with rapidly developing sequencing technologies is one of the topmost interests of various research groups [1, 8–10].

The generalizability of variant callers is an important factor [11–14] as DNA sequencing datasets exhibit diverse error profiles depending on the sequencing method. Specifically, sequencing methods are distinguished by unique combinations of protocols and experimental factors, such as the source of samples [15], sample and library preparation kits from multiple vendors [16], sequencer configurations including read length and coverage, and the type of the sequencer, all of which can result in different error profiles. Since DeepVariant [11] demonstrated deep learning (DL) based methods outperform non-DL-based methods in terms of accuracy and generalizability, much of the recent research in variant calling has focused on improving the accuracy and efficiency through introducing additional features to the input data [13, 17–22] or changing the model architecture [22–24], as well as designing a new variant calling pipeline [13, 14, 25]. However, their reliance on supervised learning poses a challenge for generalizing to sequencing methods with different error profiles, requiring large amounts of labeled data that demand expert human resources to obtain [9, 10, 26–28]. Moreover, even a subtle difference in error profiles of the sequencing data that are unseen during training, such as those due to minor changes in the sequencing methods, can challenge the robustness of DL-based variant caller (DVC) and degrade the variant calling accuracy [19, 20].

Our study introduces a new perspective, framing the challenges of the robustness and generalizability in DVC for a target sequencing method as domain adaptation and semi-supervised learning problems. We train DVC using labeled datasets and easily obtainable unlabeled datasets from target sequencing methods, each considered as a distinct domain. The labeled datasets establish our source domain, while the unlabeled or partially labeled datasets from the target sequencing method form our target domain. The generalization of DVC to a target sequencing method thus unfolds into solving two problems: (1) if only unlabeled datasets are available from the target sequencing method, it can be viewed as an unsupervised domain adaptation (UDA) problem; and (2) if partially labeled datasets are obtainable from the target sequencing method, it can be seen as a semi-supervised domain adaptation (SSDA) problem.

In this paper, we present RUN-DVC, the first semi-supervised training approach for DVC that addresses the above UDA and SSDA problems. In essence, RUN-DVC learns error profiles from unlabeled datasets of the target sequencing method using two training modules. First, RUN-DVC employs consistency training, a semi-supervised training technique, making the model generalize well on unlabeled data with unseen error profiles by propagating label information from labeled to unlabeled data. Second, RUN-DVC integrates random logit interpolation, a domain adaptation technique, aiding label propagation by reducing domain discrepancy between source and target domains that arise from varying error profiles.

We evaluate RUN-DVC in comparison with the supervised training approach on generalization scenarios using nine sequencing methods comprising 33 publicly available real-world DNA sequencing datasets [9, 10, 12]. Under UDA scenarios using short-read datasets from Illumina and BGI platforms, RUN-DVC notably increased the variant calling accuracy, enhancing SNP *F*_1_-score and INDEL *F*_1_-score by up to 6.40 %p and 9.36 %p respectively. This demonstrates that RUN-DVC improves the robustness of DVC by learning sequencing error profiles from unlabeled datasets specific to the target sequencing method. Moreover, we show the broad applicability of RUN-DVC by applying it to long-read sequencing platforms including Pacific Biosciences (PacBio) and Oxford Nanopore Technology (ONT) sequencing platforms. Finally, we demonstrate that RUN-DVC could match the variant calling accuracy of the supervised training approach using merely half of the labeled datasets in a semi-supervised domain adaptation (SSDA) scenario. This result showcases the potential of RUN-DVC to facilitate a label-efficient generalization of DVC to various sequencing methods, serving as a key advantage in practical deployment.

## Results

### Overview of RUN-DVC

We developed RUN-DVC, a semi-supervised training approach for DVCs that improves robustness and generalizability to a target sequencing method by learning error profiles from unlabeled data of the target sequencing method. RUN-DVC optimizes the DVC model through a novel loss function that combines unsupervised and supervised losses from two training modules. First, the unsupervised loss is derived from the semi-supervised learning (SSL) module that incorporates consistency training within unlabeled data. This approach uses two differently augmented versions of the same unlabeled data for training, with one serving as a model input and the model prediction on the other as a pseudo-label. By minimizing discrepancies between these, the model propagates labels from labeled data to similar unlabeled data, allowing the model to generalize well from known data to unlabeled data with different error profiles. Second, the supervised loss is derived from the random logit interpolation (RLI) module that aligns embeddings of the source and target domains. The idea is to infer the model twice every iteration with two batches: 1) a batch solely consisting of source domain data and 2) a combined batch of both source and target domain data. Subsequently, the outputs of the source domain data from both batches are interpolated and compared to the ground truth labels. This promotes the model prediction to be accurate, despite fluctuations in batch normalization layer statistics across the source and target domains, thus resulting in a model that better represents both domains.

These training modules complement the supervised training approach without changing the DVC model architecture, positioning RUN-DVC as an alternative training solution for DVCs. Fig. 1 illustrates the schematic overview of RUN-DVC.

**Fig. 1:**
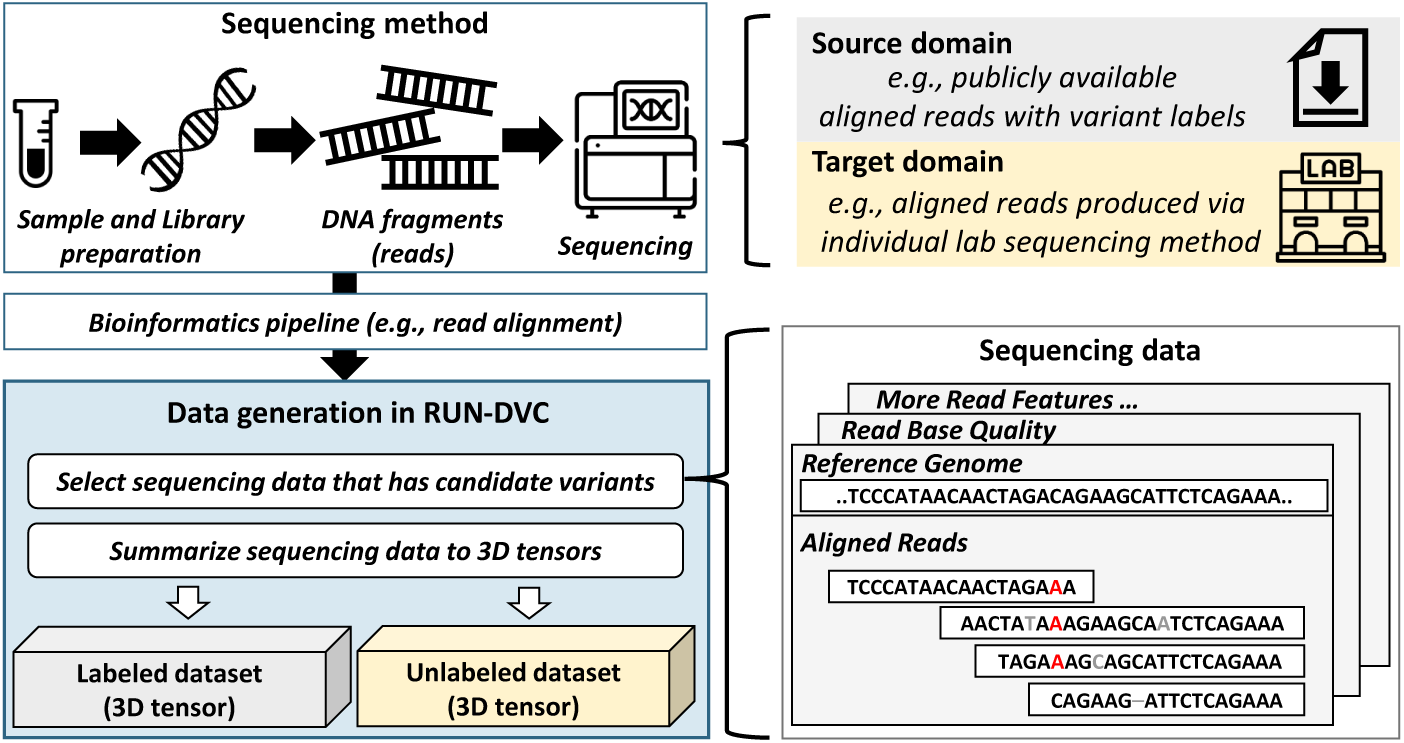
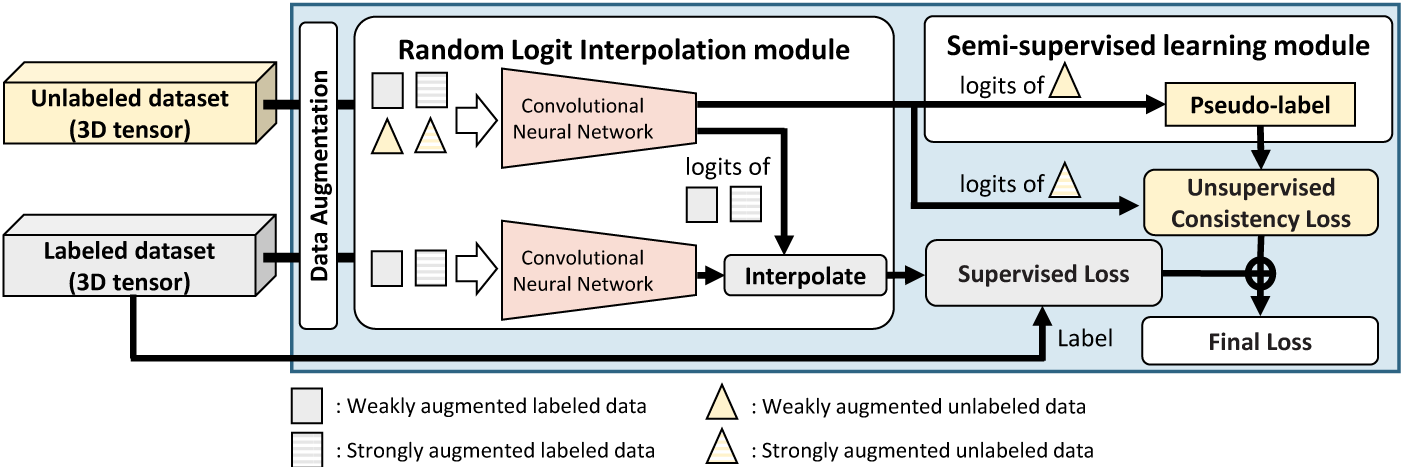
Overview of RUN-DVC workflow. (a) The overview of dataset generation for RUN-DVC. RUN-DVC addresses domain adaptation and semi-supervised learning problems, treating each sequencing method as a distinct domain. The source domain consists of labeled sequencing datasets, for example, publicly accessible sequencing data, while the target domain encompasses unlabeled or partially labeled sequencing datasets from a different sequencing method. A bioinformatics pipeline process sequencing data through stages including read alignment, sorting, and Indel realignment. Subsequently, sequencing data possessing candidate variants are selected and converted into 3-dimensional tensors. Depending on the availability of variant labels, these datasets are then categorized as either labeled (with variant labels) or unlabeled (without variant labels). (b) Illustration of the training process of RUN-DVC. RUN-DVC trains the DVC model by optimizing the sum of the supervised and unsupervised loss computed using labeled and unlabeled datasets. During each training iteration, both the random logit interpolation module and the semi-supervised learning module are used to compute the supervised and unsupervised loss, respectively. The supervised loss for labeled datasets is computed by comparing the logits obtained from the random logit interpolation module with the corresponding labels in the dataset. The unsupervised loss is computed by comparing the pseudo-label with the output of the CNN model on the strongly augmented version of the same unlabeled data. We use the class with the maximum value among predictions from the weakly augmented data as the pseudo-label. The details of the process can be found in the Methods section and the pseudocode of the training procedure is presented in the Supplementary Algorithm. S2.

### Datasets

We used 33 publicly available sequencing datasets from GIAB [8, 9, 26, 27] (Genome in a Bottle), the Human Pangenome Reference Consortium [29], and Google [10]. We leveraged the version 4.2.1 GIAB truth variant sets [28] as our ground-truth label for analysis. Sequencing datasets include human samples, NA12878/HG001 from 1000 Genomes [30], and two trios (HG002-HG003-HG004 and HG005-HG006-HG007) from participants in the Personal Genomes Project [31]. The summary of the sequencing datasets used in experiments is organized in Table 1 and its corresponding web links to details of sequencing methods are organized in Supplementary Table. S1.

**Table 1:** Overview of datasets used. Dataset identifier, sequencing method, number of data, and human sample numbers of sequencing datasets are provided. Note that P-A consists of two sequencing datasets from the same sample. The sample number corresponds to the suffix of the sample name (e.g., 1 stands for HG001).

| Identifier | Sequencing method (number of data) | Sample number |
| --- | --- | --- |
| I-Source | Illumina NovaSeq PCR-free 30x data (20 million) | 5,6,7 |
| I-A | Illumina NovaSeq PCR-positive 30x data (23 million) | 1,2,3,4 |
| I-B | Illumina Hiseq2500x 21-30x data (86 million) | 1,2,3,4 |
| I-C | BGISEQ-500 40x data (21 million) | 1,2,3,4 |
| I-D | Illumina HiseqX PCR-positive 30x data (20 million) | 1,2,3,4 |
| P-Source | PacBio HiFi Sequel II 30x and 50x data (26 million) | 1,2,5,5* |
| P-A | PacBio HiFi Sequel I 30x data (15 million) | 2,2* |
| O-Source | ONT SUP mode Guppy v5.0.14 50x data (34 million) | 2,3,4,5 |
| O-A | ONT HAC mode Guppy v5.0.14 50x data (34 million) | 2,3,4,5 |

Each of the datasets I-A, I-B, I-C, and I-D, exhibits distinct error profiles attributable to the different sequencing methods employed. Specifically, I-A and I-Source were sequenced using the same machine (Illumina NovaSeq) in the same institution, but I-A utilized PCR amplification in the library preparation step. I-D, processed in the same institution as I-Source, employed PCR amplification in the library preparation step and used a different sequencing machine (Illumina HiseqX). Furthermore, I-B was generated by 10x Genomics and sequenced on the Illumina Hiseq2500x platform, while I-C was produced by BGI and sequenced using the BGISEQ-500 machine. Turning to the PacBio HiFi datasets, P-Source and P-A were obtained from the Sequel II and Sequel I systems, respectively. Finally, O-Source and O-A datasets are ONT sequencing datasets, with base calls made via super accuracy (SUP) and high accuracy (HAC) modes of Guppy 5.0.14 on the ONT PromethION platform, respectively.

### Baseline methods

*BaselineBN* [32] represents a supervised training strategy employed in exist-ing DVCs supplemented with a minimal domain adaptation technique. This approach trains on labeled datasets from the source domain and also utilizes unlabeled datasets from the target domain to update batch norm statistics, fostering domain alignment[33, 34]. Another approach, referred to as *Full-label*, illustrates the maximum accuracy attainable by the model when trained on fully labeled datasets from the target domain. Data augmentation techniques are utilized during training for both *BaselineBN* and *Full-label*.

### Performance on short-read sequencing platforms

We compare the variant calling performance of RUN-DVC against *BaselineBN* and *Full-label* under four UDA settings using short-read datasets. Specifically, we train RUN-DVC and *BaselineBN* using labeled datasets from I-Source as the source domain and unlabeled datasets from I-A, I-B, I-C, and I-D as the target domains. The *Full-label* is trained using labeled datasets specific to each of I-A, I-B, I-C, and I-D. For the purposes of evaluation, the HG003 sample is excluded from all training datasets. In addition, we provide the accuracy of state-of-the-art methods, Clair3 v1.0.0 [14] and DeepVariant v1.5.0 [11], for validation of the problem and implementation. DeepVariant uses 52 sequencing datasets from various sequencing methods (including I-Source, I-A, and I-D) that account for 815,200,320 training samples, whereas Clair3 uses 12 PCR-free sequencing datasets (comprising I-Source dataset).

Fig. 2(a) shows the precision-recall curve and the best *F*_1_ scores achieved by selecting the optimal quality score threshold for each method. The Precision, Recall, and *F*_1_ score of variant calling by each method with default setting can be found in Supplementary Table. S2 and Table. S3. RUN-DVC demonstrated superior performance over *BaselineBN* across all datasets, utilizing only unlabeled datasets from the target domain. For SNP calling, the performance of *BaselineBN* is only slightly lower than *Full-label* on the I-A, I-C, and I-D datasets. However, there was a significant decrease of 7.91 percentage points (%p) in the SNP *F*_1_ score on the I-B dataset. On the other hand, RUN-DVC performed better than *BaselineBN* by improving the score by 6.40 %p, thus reducing the performance drop compared to *Full-label* to only 1.51 %p. Regarding INDEL calling, *BaselineBN* exhibited substantial performance degradation across all datasets compared to *Full-label*. However, RUN-DVC significantly reduced the performance disparity between *BaselineBN* and *Full-label*, outpacing *BaselineBN* by 2.94 %p, 9.36 %p, 1.69 %p, and 4.21 %p on the I-A, I-B, I-C, and I-D datasets, respectively. These results highlight the ability of RUN-DVC to enhance the robustness of DVC, thereby enhancing variant calling performance in the target sequencing method.

**Fig. 2:**
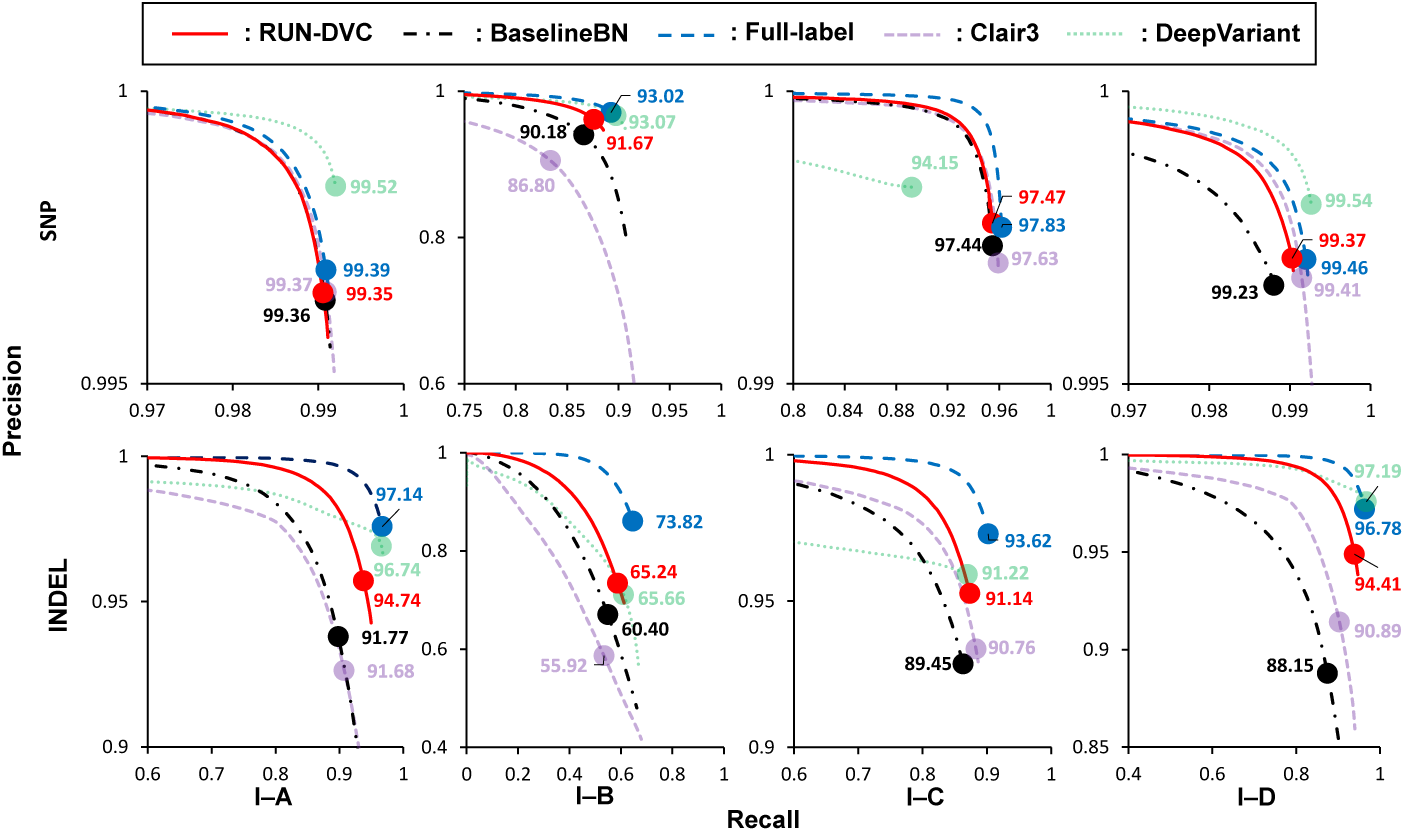
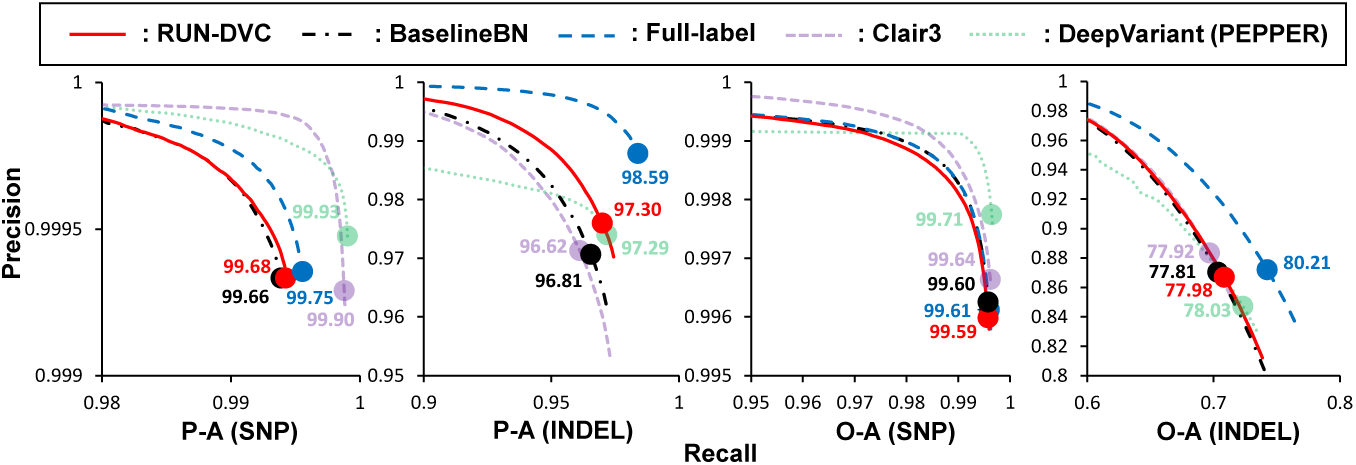
Performance of RUN-DVC under UDA setting. The quality scores are used to make precision-recall curves. The highest *F*_1_-score (percentage) achieved by each method is marked with a circle. The precision, recall, and *F*_1_ score of PASS calls are available in supplementary Table. S2, Table. S3, and Table. S4. (a) Precision-recall analysis on the HG003 sample of I-A, I-B, I-C, and I-D datasets. (b) Precision-recall analysis on the HG002 sample of the P-A dataset and the HG003 sample of the O-A dataset.

State-of-the-art DVCs, such as Clair3 and DeepVariant, displayed discernible performance reductions on certain datasets. Specifically, DeepVariant’s performance faltered on I-B and I-C datasets, while Clair3’s performance diminished across all datasets. DeepVariant showed no performance degradation in SNP and INDEL calling on the I-A and I-D datasets, which are part of the training dataset of DeepVariant. However, relative to the *Full-label* method, it registered a decrease in the SNP F1 score by 3.68 %p on I-C and a decrease in the INDEL F1 score by 12.90 %p and 2.37 %p on I-B and I-C datasets, respectively. In the case of Clair3, it experienced a 21.99 %p decrease in the SNP *F*_1_ score on I-B and a decline in INDEL *F*_1_ scores by 6.43 %p, 22.16 %p, 2.86 %p, and 6.97 %p on the I-A, I-B, I-C, and I-D datasets, relative to *Full-label*. These observations demonstrate that state-of-the-art DVCs may struggle to maintain robustness when confronted with sequencing methods unseen during training.

### Performance on long-read sequencing platforms

In order to demonstrate the broad applicability of RUN-DVC across various sequencing methods, we compared its performance with that of *BaselineBN* on two UDA settings from each PacBio and ONT dataset. Within these scenarios, P-Source and O-Source were designated as the source domains, while P-A and O-A constituted their respective target domains. Due to the limited availability of publicly accessible PacBio Sequel I datasets generated with the same sequencing method, the HG002 sample’s unlabeled datasets were used during the training phase, and the same sample was employed for evaluation. We provide the variant calling accuracy of models provided by Clair3 and DeepVariant (PEPPER) for PacBio Sequel II and ONT Guppy5 SUP mode datasets. Details of training datasets used for Clair3 and DeepVariant can be found in Supplementary Note. 4.

Fig. 2(b) shows the results. For both PacBio and ONT datasets, RUN-DVC incorporated haplotype information as an additional feature within the input. Furthermore, for the ONT datasets, RUN-DVC employed an input tensor of a different size, supporting a read depth of up to 89 as opposed to a read depth of 55 utilized for PacBio and short-read datasets (refer to Supplementary Note 1 for more details on input tensor). Nevertheless, RUN-DVC outperformed *BaselineBN* on P-A and O-A datasets even when using the same set of hyperparameters that was used in short-read datasets. These results confirm the versatility of RUN-DVC across diverse sequencing methods ranging from short reads to long reads that use different input sizes and input features.confirm the versatility of RUN-DVC across diverse sequencing methods ranging from short reads to long reads that use different input sizes and input features.

### RUN-DVC effectively learns sequencing error profiles from unlabeled datasets

To ascertain whether the observed performance improvement stemmed from the learning of error profiles of the target sequencing method, we conducted an additional analysis. This involved contrasting the performance of RUN-DVC with both *BaselineBN* and *Full-label* in relation to genomic contexts. RUN-DVC and *BaselineBN* were trained under two UDA settings using labeled datasets from I-Source as the source domain and unlabeled datasets (excluding the HG003 sample) from I-A or I-B as the target domain. We report the variant-calling performance in the genome’s difficult-to-map regions (low-mappability and segmental duplications regions) and low-complexity regions (tandem repeats and homopolymer regions) according to the GIAB v2.0 stratification data.

Fig. 3(a) shows the variant calling performance on the HG003 sample of the I-A dataset. *BaselineBN* exhibited notable performance degradation in INDEL calling accuracy only in the low-complexity regions. Specifically, in each tandem repeats region and homopolymers region, RUN-DVC achieved *F*_1_-score 0.9374 and 0.8875 which are 1.07 %p and 5.64 %p higher compared to *BaselineBN* that achieved 0.9267 and 0.8311. For whole regions except for low-complexity regions, RUN-DVC (SNP *F*_1_-score: 0.9937, INDEL *F*_1_-score: 0.9931), *BaselineBN* (SNP *F*_1_-score: 0.9938, INDEL *F*_1_-score: 0.9930), *Full-label* (SNP *F*_1_-score: 0.9939, INDEL *F*_1_-score: 0.9932) achieved similar SNP and INDEL *F*_1_-scores. These outcomes align with the known impact of PCR amplifications, which introduce various artifacts on repetitive DNAs [35]. We conjecture that the absence of these artifacts in the PCR-free I-Source dataset is the reason for the *BaselineBN* ’s performance degradation in the tandem repeats and homopolymers regions. Notably, RUN-DVC effectively counteracts this effect, improving variant calling accuracy and confirming its ability to learn error profiles, including artifacts and deletions caused by PCR amplification, from unlabeled datasets.

**Fig. 3:**
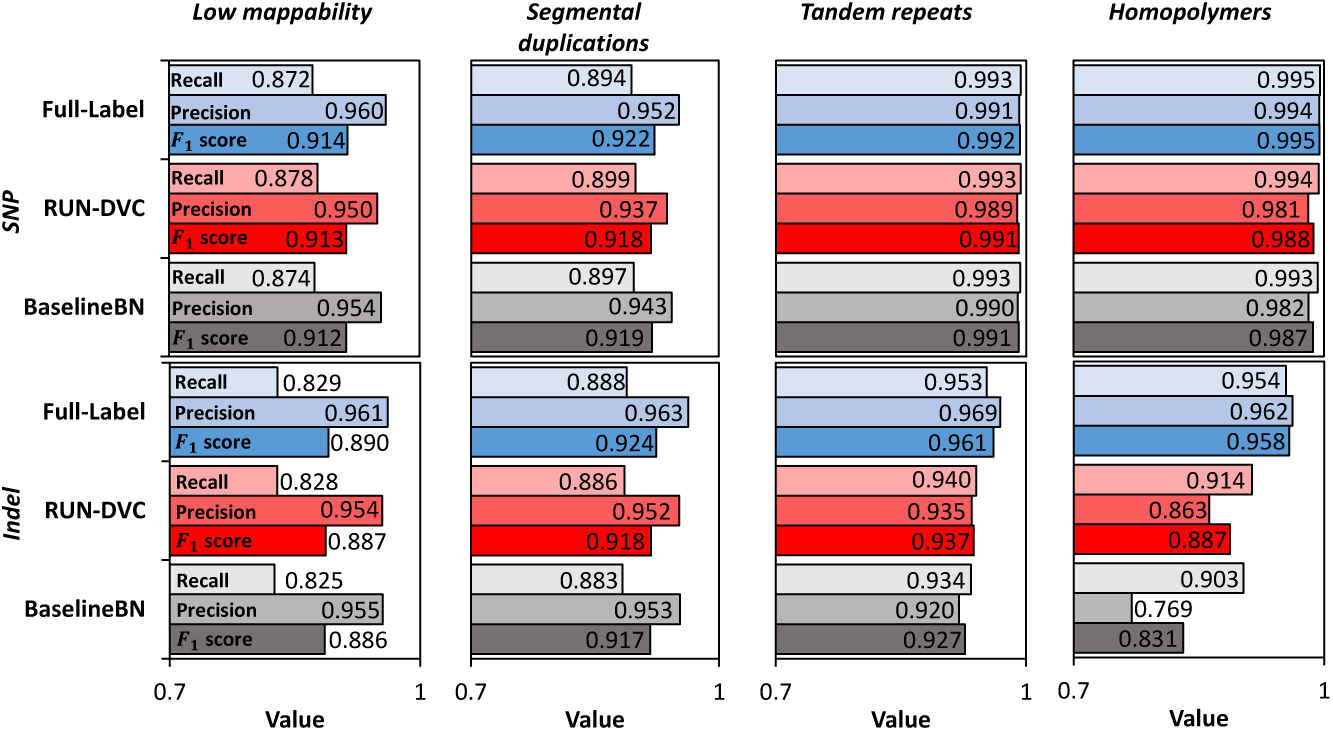
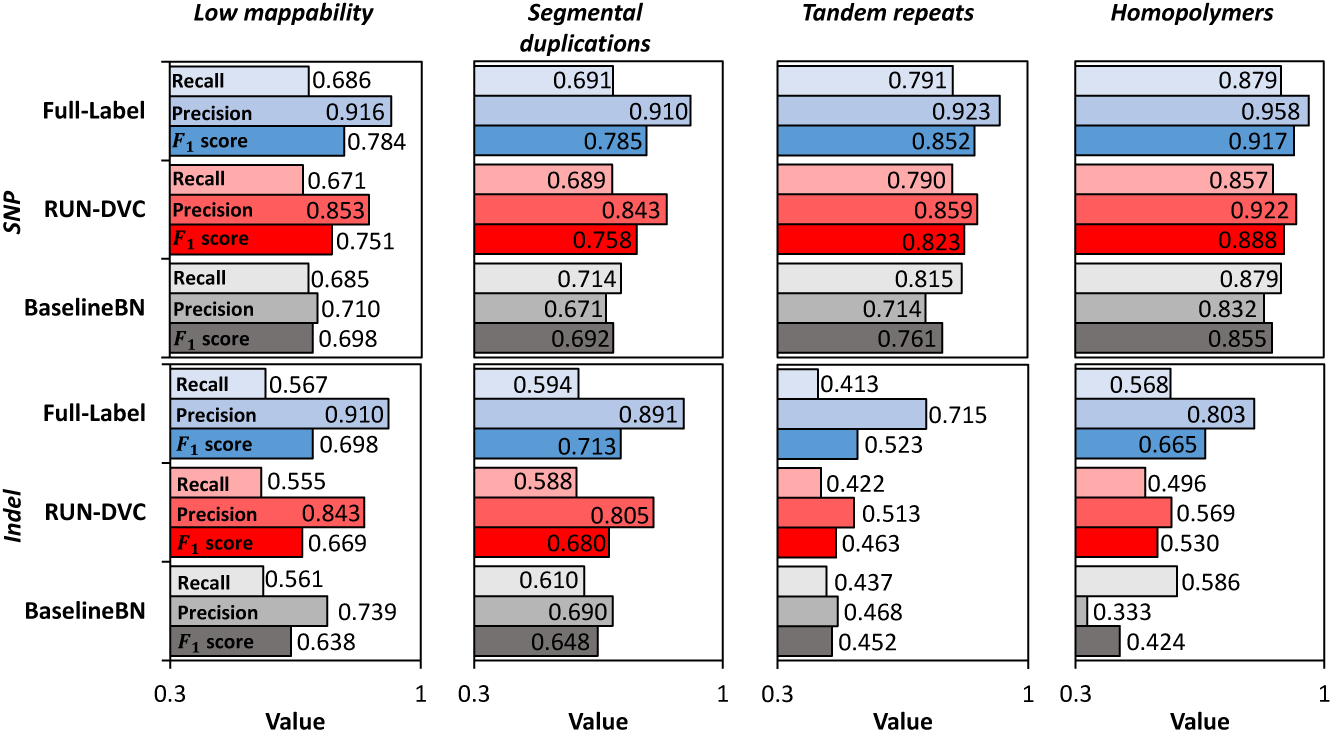
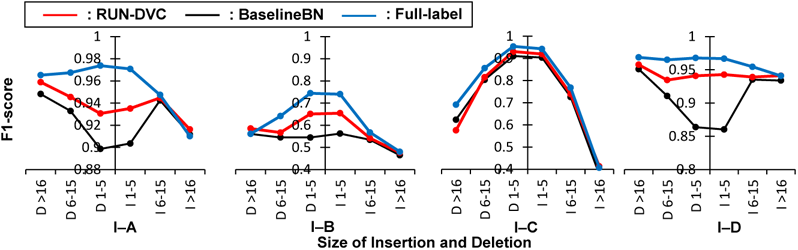
Performance analysis of RUN-DVC. (a) Performance analysis by genomic regions on HG003 sample of I-A dataset. (b) Performance analysis by genomic regions on HG003 sample of I-B dataset. (c) Performance analysis for different INDEL sizes on HG003 sample. (x-axis represents the size of INDELs, I: insertion, D: deletion)

Fig. 3(b) shows the variant calling performance on the HG003 sample of the I-B dataset. RUN-DVC significantly outperforms *BaselineBN* in both SNP and INDEL calling accuracy across all regions. Particularly in regions of low mappability, segmental duplications, tandem repeats, and homopolymers, RUN-DVC surpasses *BaselineBN* with SNP *F*_1_-scores improved by 5.335 %p, 6.619 %p, 6.141 %p, and 3.334 %p, and INDEL *F*_1_ scores by 3.134 %p, 3.208 %p, 1.142 %p, and 10.59 %p respectively. In our analysis, the I-B dataset emerged as the most demanding scenario for generalization, exhibiting the most pronounced performance deterioration for all *BaselineBN*, DeepVariant, and Clair3. This is attributed to its uniquely disparate error profiles and the extensive magnitude of errors dispersed throughout the entire genomic regions. Despite this, RUN-DVC excelled, demonstrating its capacity to learn and adapt to significantly different error profiles even in a demanding variant calling case.

Finally, we assess the INDEL calling performance of RUN-DVC for different INDEL sizes as shown in Fig. 3(c). RUN-DVC consistently surpasses the performance of *BaselineBN*, underscoring its efficacy for a range of INDEL sizes. In summary, RUN-DVC’s strength stems from its ability to learn a multitude of error profiles from unlabeled datasets, showcasing versatility that is not confined to specific regions or types of variants.

### Ablation study

To elucidate the individual contributions of the two training modules of RUN-DVC to the overall performance, we perform an ablation study. In this experiment, we compare *BaselineBN*, “RUN-DVC w/o RLI” that was trained solely via the semi-supervised learning module without the use of random logit interpolation module, and the full RUN-DVC. All methods are trained on two UDA settings using labeled datasets from I-Source and unlabeled datasets (excluding the HG003 sample) from I-A or I-D datasets.

Fig. 4(a) shows the results of RUN-DVC, “RUN-DVC w/o RLI“, and *BaselineBN* on HG003 sample of I-A and I-D datasets over 4 independent runs. In the case of the I-A dataset, RUN-DVC delivered the highest accuracy (SNP *F*_1_-score: 0.9935, INDEL *F*_1_-score: 0.9387), outperforming “RUN-DVC w/o RLI” (SNP *F*_1_-score: 0.9934, INDEL *F*_1_-score: 0.9243) and *BaselineBN* (SNP *F*_1_-score: 0.9934, INDEL *F*_1_-score: 0.9069). For the I-D dataset, while RUN-DVC demonstrated superior accuracy (SNP *F*_1_-score: 0.9938, INDEL *F*_1_-score: 0.9413) as compared to *BaselineBN* (SNP *F*_1_-score: 0.9927, INDEL *F*_1_-score: 0.8697), “RUN-DVC w/o RLI” exhibited a downturn in SNP calling accuracy (SNP *F*_1_-score: 0.9290, INDEL *F*_1_-score: 0.9181). Furthermore, the variant calling performance on the I-D dataset by “RUN-DVC w/o RLI” displayed remarkable instability.

**Fig. 4:**
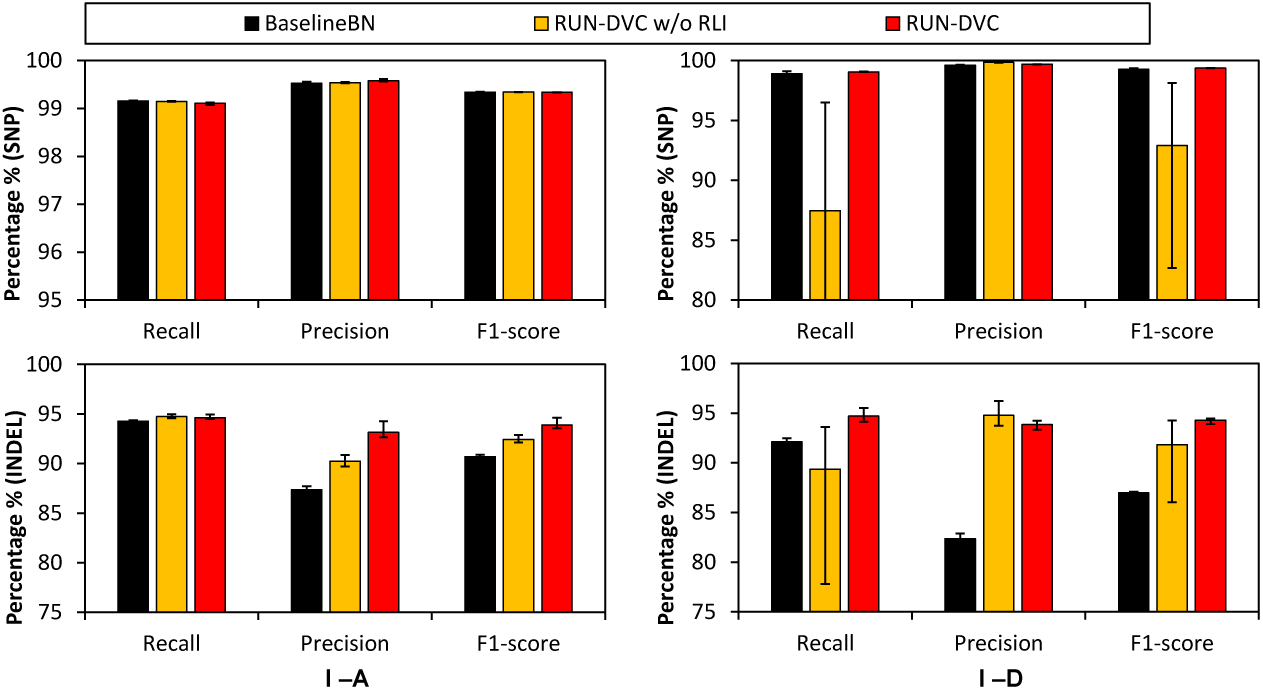
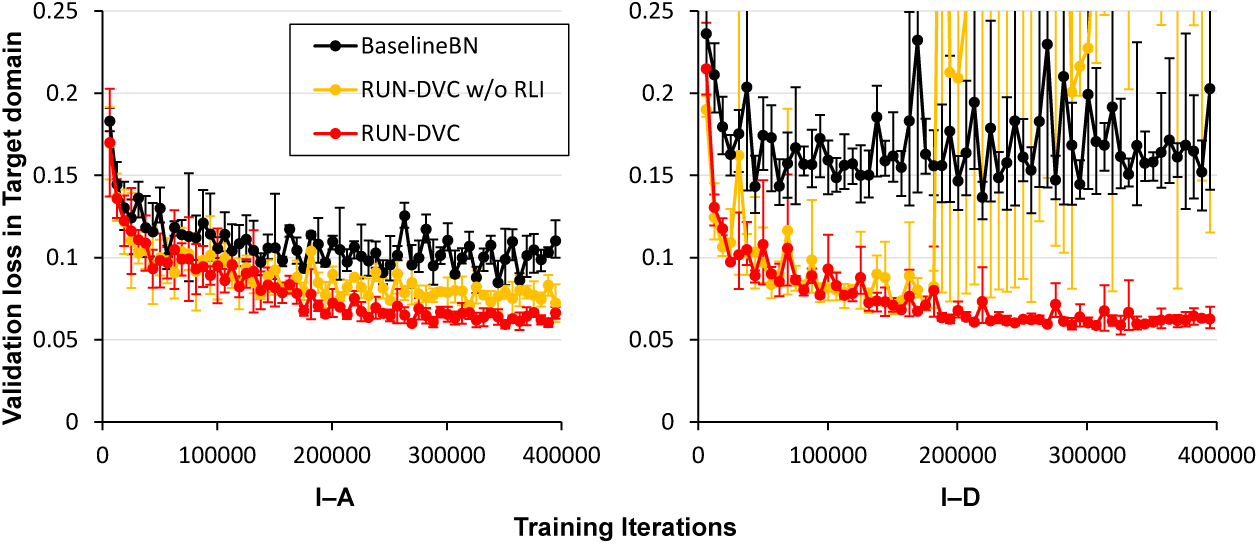
Ablation study. Error bars represent the range between the maximum and minimum values. (a) Performance of *BaselineBN*, “RUN-DVC w/o RLI“, and RUN-DVC in HG003 sample of I-A and I-D dataset. (b) Validation loss curve of *BaselineBN*, “RUN-DVC w/o RLI“, and RUN-DVC during training.

Fig. 4(b) shows the validation loss on the target domain over the course of training iterations for *BaselineBN*, “RUN-DVC w/o RLI“, and RUN-DVC. Remarkably, the validation loss of “RUN-DVC w/o RLI” on the I-D dataset exhibited substantial fluctuations and failed to converge to a favorable solution, unlike RUN-DVC. This suggests the crucial role of the RLI module in aligning domains and ensuring stable learning on the target domain.

Overall, these observations underscore the vital role of the SSL and RLI modules in the efficacy of RUN-DVC. It is also pertinent to note that, we proposed multiple data augmentation strategies for DNA sequencing datasets that are also crucial for the overall performance of RUN-DVC. Further details and analysis on these data augmentation strategies can be found in Supplementary Fig. S3.

### Using RUN-DVC for foreseeing when DVC fails to generalize

To demonstrate the utility of RUN-DVC as an indicator for generalization failure of DVC in a target sequencing method, we examined the variant calling outcomes of RUN-DVC and the supervised trained model. This exploration entails analyzing disparities in quantities, focusing on both biallelic changes in SNPs and INDELs of varying sizes, and comparing these between RUN-DVC and *BaselineBN* under the UDA setting. The results are shown in Supplementary Fig. S2.

Our evaluation of the number of biallelic base alterations offers meaningful insights into the potential generalization failure of *BaselineBN* in relation to SNP identification performance. Notably, we did not observe any significant discrepancy in the number of biallelic base alterations in the I-A, I-C, and I-D datasets. However, a noticeable difference surfaced in the I-B dataset, in which a large performance degradation in SNP calling of *BaselineBN* was observed.

As for INDEL calling accuracy, the number of INDEL variants by size serves as a key indicator of potential generalization failures in *BaselineBN*. Across the I-A, I-B, I-C, and I-D datasets, the total number of disparities for each INDEL size between RUN-DVC and *BaselineBN* was 34,528, 222,193, 15,497, and 47,324 respectively, which seems to correlate with the degree of performance degradation.

In addition, we also assess the distribution of genotypes as shown in Supplementary Fig. S1(b). Intriguingly, we observe that the variant calling quality scores of *BaselineBN* could either exceed or fall short of those from RUN-DVC and *Full-label*, despite a considerable decrease in variant calling accuracy. Therefore, in light of these findings, we suggest that a comparison of both the quantities of biallelic base changes and the numbers of INDELs of varying sizes, produced by RUN-DVC and the model trained through the supervised method, could serve as effective indicators for instances of generalization failure of DVCs.

### RUN-DVC enables label-efficient generalization of DVC to target sequencing methods

To assess the generalizability of RUN-DVC, we conducted a comparison of the validation loss and variant calling accuracy between RUN-DVC and *BaselineBN* under varying quantities of target domain labeled datasets. The results over 2 independent runs are shown in Fig. 5 (the numbers can be found in Supplementary Table. S7). This evaluation was conducted within two SSDA scenarios, utilizing I-Source as the source domain and I-A and I-B as target domains. The validation loss was determined by evaluating the performance of both methods against identical validation datasets, comprising 5% of the total target domain datasets that were excluded from the training phase. The variant calling accuracy was measured on the HG003 sample that was excluded from the training phase. Importantly, RUN-DVC consistently surpassed the variant calling performance of *BaselineBN* given the same amount of labeled datasets. Additionally, in terms of validation loss, RUN-DVC exhibited almost double the label-efficiency compared to *BaselineBN*. These findings underscore the superior generalizability of RUN-DVC, manifested by its enhanced ability to effectively propagate label information from the labeled datasets to unlabeled datasets.

**Fig. 5:**
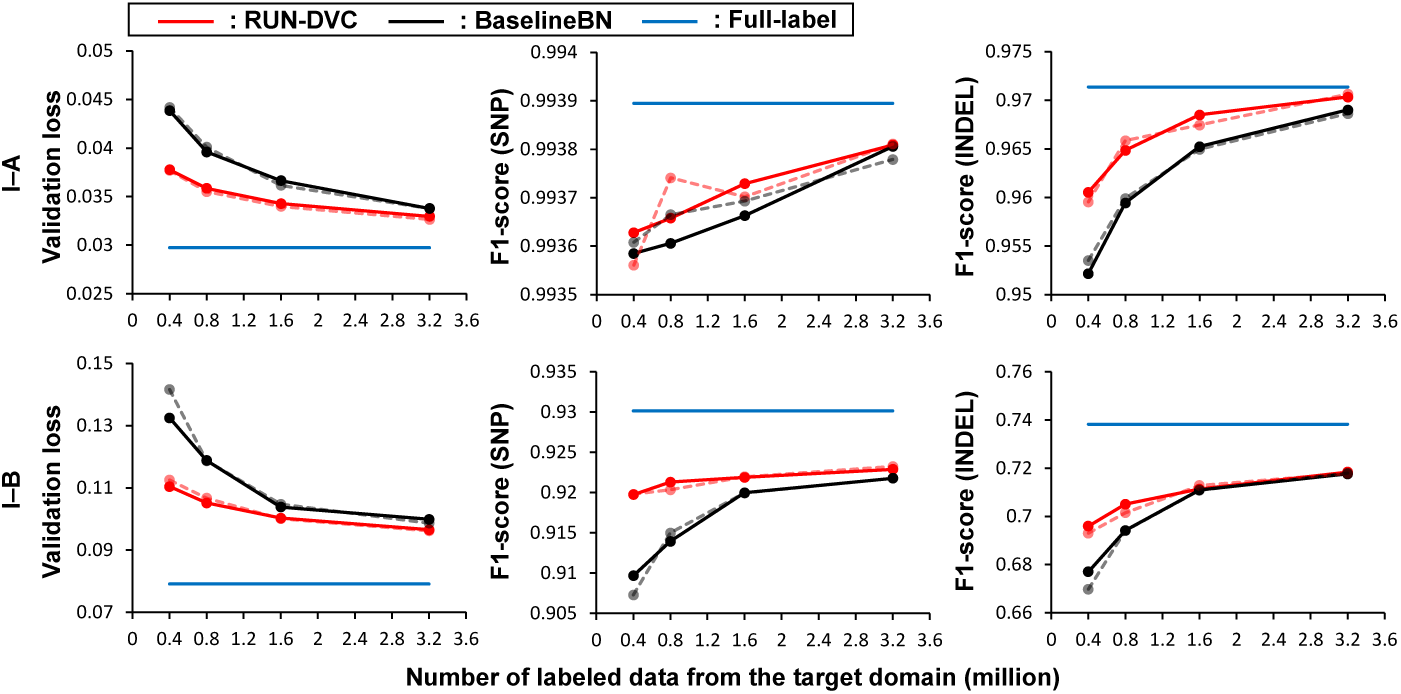
Performance of RUN-DVC under various quantities of labeled datasets in SSDA setting. Transparent lines are the results using a different random seed for target domain labeled datasets selection.

## Discussion

We present RUN-DVC, the first semi-supervised training approach for DVC that improves the robustness and enables label-efficient generalization to a target sequencing method. RUN-DVC leverages the consistency training and random logit interpolation techniques, allowing it to learn error profiles from unlabeled data of the target sequencing method using the knowledge obtained in labeled data. These training techniques are complementary to the supervised training approach, positioning RUN-DVC as an alternative training solution for existing DVCs.

In the UDA experimental setup, RUN-DVC exceeds the performance of the supervised training approach in the accuracy of variant calling. This signifies the potential of RUN-DVC to enhance the robustness of DVC against specific sequencing methods by learning error profiles in unlabeled datasets. Furthermore, we have established the applicability of RUN-DVC to long-read sequencing platforms such as PacBio and ONT. This demonstrates the extensive adaptability of RUN-DVC to diverse sequencing methods, encompassing both short reads and long reads. Looking forward, we believe that incorporating RUN-DVC into existing DVCs [11, 17–24, 36] will expand their utilization across a variety of sequencing methods, leveraging the potential of unlabeled data.

One of the prominent applications of RUN-DVC is the generalization of DVC across various species. RUN-DVC operates without making specific assumptions regarding the sequencing method, including the species. This makes it possible to apply RUN-DVC to sequencing data from species other than humans. However, our evaluation was constrained by the limited availability of sequencing datasets using the same method, and the absence of benchmark datasets, preventing a comprehensive assessment in this regard.

We have demonstrated that RUN-DVC can be used to foresee the generalization failure of DVC by examining numbers and types of variants produced by both RUN-DVC and the supervised trained model. This can be used to determine whether further training of the DVC model is required to achieve a higher variant calling accuracy in the target sequencing method.

Finally, RUN-DVC outperforms the supervised training method by achieving higher variant calling accuracy with fewer labeled datasets, thus illustrating that RUN-DVC offers a more label-efficient training solution. For individual laboratories employing custom sequencing methods and seeking to establish an accurate variant calling pipeline with DVC, RUN-DVC is expected to be particularly valuable, especially when combined with the retraining solution [37].

We acknowledge several limitations of RUN-DVC. First, the RLI module is applicable only for DVCs that use a CNN model with batch normalization layers, a characteristic that is true for the majority of existing DVCs [11, 13, 14, 17, 19, 23, 24]. Nevertheless, shifting to a different model architecture other than CNN might necessitate the development of new methods to replace the RLI module. Second, we have provided evidence that RUN-DVC, operating on a single hyperparameter setting, is effective across a range of sequencing datasets, outperforming the supervised training method. However, we acknowledge that the application of optimal hyperparameters and data augmentations might drive further enhancements in RUN-DVC’s performance (see Supplementary Fig. S3 for an ablation study on data augmentations). A third limitation lies within the SSDA setting where the target domain’s labeled datasets selection is randomized. We postulate that the integration of active learning strategies could potentially augment RUN-DVC’s label efficiency. Lastly, RUN-DVC has yet to be validated for somatic mutation calling—an application with different task requirements and fundamental assumptions. Future investigations should address this by extending the use of RUN-DVC to include somatic mutation callers, particularly in cases where the sequencing datasets exhibit a greater degree of diversity.

## Methods

### Evaluation metrics

We used Illumina’s Haplotype Comparison Tools [12] (hap.py) and GIAB v4.2.1 truth variant data to benchmark variant-calling results. Hap.py generates three metrics Precision, Recall, and F1-score for each of the categories SNP and Indel, respectively. From the number of true positives (TP), false positives (FP), true negatives (TN), and false negatives (FN), hap.py computes the three metrics as 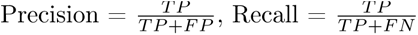, and 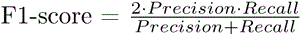.

### Input/output and model architecture

#### Selecting candidate variants

RUN-DVC selects candidate variants for training data or variant calling using the following algorithm. For each position in the reference genome, all the reads that overlap the position are collected. Next, RUN-DVC selects a position as a candidate variant if two conditions are met: (1) the number of aligned reads surpasses the coverage threshold. (2) the percentage of mismatches between the reference genome and the aligned reads is above the allele frequency threshold.

#### Input tensor

RUN-DVC takes a 3-dimensional tensor as input for the DNN model, which is structured similarly to an image. For instance, in the case of short reads, the input tensor has a shape of 55 rows, 33 columns, and 7 channels. Specifically, a 3-dimensional tensor comprises information about a candidate variant region. Each channel in the input tensor represents a unique feature of the sequencing data including reference bases, variants in reads, strand information, mapping quality, base quality, the proportion of candidate variants, insertion bases, and phasing information. The columns correspond to positions in the reference genome, while the rows represent individual reads aligned to those positions. A detailed explanation of each channel is provided in Supplementary Note. 1, and an example of an input tensor is shown in Supplementary Fig. S5.

#### Output of the DNN model

During inference, the DNN model performs four classification tasks on the input. These tasks involve predicting four variables: *G*, *Z*, *L*_1_, and *L*_2_, where *G* corresponds to 21 possible genotypes and *Z* represents the zygosity of the variant. The variables *L*_1_ and *L*_2_ indicate the length of the two variants in the diploid organism, with possible values ranging from -16 to 16. Note that, the values of -16 and 16 for *L*_1_ and *L*_2_ represent variants with lengths larger than 16. Specifically, the set of possible values for each variable is as follows:

- *G ∈*{ AA, AC, AG, AT, AIns, ADel, CC, CG, …, DelDel}
- *Z ∈* {Homozygous reference, Heterozygous, Homozygous non-reference}
- *L*_1_*, L*_2_ *∈*{*−*16*, …,* 16}

#### Constructing labeled training dataset

RUN-DVC constructs a labeled training dataset from the candidate variants using ground truth labels. Specifically, each data point is selected from candidate variants, which consists of pairs of a tensor that summarizes a candidate variant and its corresponding variant label that includes four labels *G*, *Z*, *L*_1_, and *L*_2_. *N_m_* true variants and *N_r_*non-variants are selected from candidate variants, in a ratio specified by the target ratio 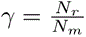. RUN-DVC constructs labeled training dataset using *γ* = 1 [14].

#### Constructing unlabeled training dataset

Different sequencing methods can generate candidate variants with vastly different true variants to non-variants ratios. For instance, the ONT dataset may have 100 times more non-variants than true variants, while the Illumina dataset has a similar number of non-variants and true variants. This results in the ONT dataset having 50 times longer training iterations and sizes compared to the Illumina dataset. However, it has been shown including non-variants beyond a certain number offers marginal improvement in performance [14].

To reduce the number of non-variants in the unlabeled dataset, an RNN model trained on the labeled dataset is used to obtain *confidence scores* for candidate variants that are probabilities of being true variants. The RNN model consists of two bidirectional long short-term memory (Bi-LSTM) layers with 128 and 160 LSTM units, and uses the pileup input proposed by Clair3 (see Supplementary Note. 1 for details of pileup input). We observed the RNN model shows high accuracy in filtering non-variants from candidate variants (see Supplementary Table. S5 and Table. S6 for details). Filtering out non-variants reduces the size of the unlabeled dataset while still including the true variants necessary for training.

The unlabeled training dataset is constructed using all candidate variants when the total number of candidate variants in a single sample is below a threshold value of *T* . However, if the total number of candidate variants exceeds the threshold, the top *T* candidate variants are selected based on the *confidence scores*. In our experiments, 80 million is used for the threshold value *T* . Our approach ensures that the training dataset contains enough true variants for effective training while reducing the size of the dataset and training time by removing non-variants.

#### Model architecture

RUN-DVC trains a convolutional neural network (CNN) model on 3-dimensional input tensors that each comprises information about a candidate variant region. The CNN model comprises three blocks, each consisting of a basic convolution block followed by a standard residual block. After the blocks, an adaptive max pooling layer is used at the end of the blocks that outputs 512 features. The basic convolution block includes a convolutional block, a batch normalization layer, and a Leaky Relu activation layer, which decreases the width and height of features. The classifier head comprises two dense layers for each task that output 256 and 128 features, respectively. The architecture of the CNN model is presented in Supplementary Fig. S4.

### Training objectives of RUN-DVC

RUN-DVC solves the unsupervised and semi-supervised domain adaptation problem by leveraging the labeled dataset from the source domain and the unlabeled dataset from the target domain. In the case of the SSDA setting, the labeled samples from the target domain are included in the labeled dataset. RUN-DVC minimizes the loss function *ℒ* that consists of two losses: (1) a supervised classification loss *ℒ_l_* on the labeled sample, (2) an unsupervised classification loss *ℒ_u_* on the unlabeled sample. The DNN model in RUN-DVC performs four classification tasks, hence, the loss function to be optimized is expressed as:

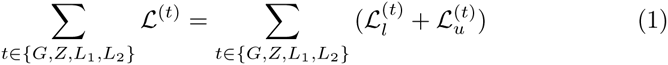

In essence, RUN-DVC optimizes the sum of both supervised and unsupervised loss functions to solve unsupervised and semi-supervised domain adaptation problems. We provide the pseudocode of the training procedure in Supplementary Algorithm. S2.

#### Weak and strong data augmentations

RUN-DVC adapts the concept of weak and strong augmentation from computer vision. In computer vision, weak augmentation comprises augmentations such as flipping and rotating, whereas strong augmentation includes complex transformations such as changes in color and image structure. For RUN-DVC, weak augmentations consist of subsampling and vertical shifting, while strong augmentations additionally employ feature distortions. During training, the augmentations are applied to batches of data from both labeled and unlabeled datasets at each iteration. This is in contrast to Clair3 and DeepVariant, where augmentations are applied before constructing the training dataset. Applying augmentations during training enables the same data to be augmented in a different way during each epoch of training, which enhances the model’s resilience against variability. Further details regarding weak and strong augmentations can be found in the *Data Augmentations* subsection.

#### Notation

We represent a batch of labeled samples from the source domain by *B_l_*, and a batch of unlabeled examples from the target domain by *B_u_*. Each batch is composed of samples that are weakly and strongly augmented, which we denote respectively by *B_l,weak_*, *B_l,strong_*, *B_u,weak_*, and *B_u,strong_*.

#### Supervised classification loss is obtained using random logit interpo-lation

RUN-DVC addresses the domain adaptation problem by using random logit interpolation [32] (RLI) during training. RLI interpolates the logits (the outputs of the last layer in the DNN model before the activation layer) from both source and target domains to generate more representative batch statistics in batch normalization layers. During each iteration of training, the DNN model is inferred twice to obtain two logits, *O_c_* and *O_l_*. *O_c_* is computed using a batch *{B_l_, B_u_}* that includes both labeled and unlabeled datasets, whereas *O_l_* is computed using a batch *{B_l_}* that only contains the labeled dataset:

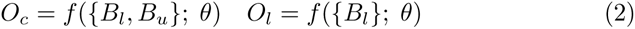

Importantly, the batch normalization layers in the DNN model are updated during the computation of *O_c_*, whereas they remain fixed during the computation of *O_l_*. This ensures that model parameters and batch normalization layers are adapted to the target domain during training.

The supervised classification loss *ℒ_l_*is computed using cross entropy loss between ground truth label *Y* and the final logits *O_RLI_*, where the final logits *O_RLI_*for *B_l_* labeled samples are obtained by randomly interpolating *O_c_* and *O_l_*, as shown below:

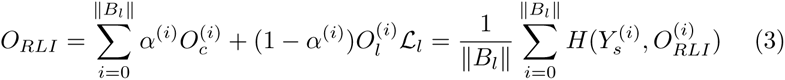

where *α ∈* ℝ*^Bl×k^* is a vector of random values drawn from uniform distri-bution *U ^Bl∗k^* (0, 1), function *f* denotes the DNN model that outputs *k* logits for each sample, and *H*(*Y, P*) denotes the cross-entropy between label *Y* and logits *P* .

#### Unsupervised classification loss is obtained via semi-supervised learn-ing

The semi-supervised learning module employs the consistency training technique [38] to learn from the unlabeled datasets. At each iteration, the model is trained to minimize the loss between the pseudo-labels generated from the weakly augmented data and the output generated from the strongly augmented data. This improves the model’s robustness against input data variability as the outputs of weakly and strongly augmented data are encouraged to be consistent. Also, the label information is gradually propagated from the labeled data to the unlabeled data [39].

The unlabeled samples’ logits *O_u_* is obtained from the logits *O_c_* of combined batches as follows:

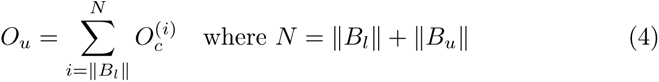

where *O_u_*contains the logits of both weakly and strongly augmented unlabeled samples, denoted as *O_u,weak_*and *O_u,strong_*, respectively:

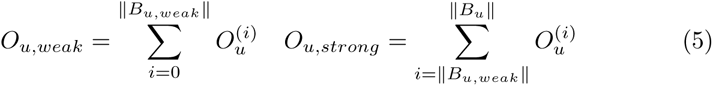

To prevent the model from training with inaccurate pseudo-labels during the early stages of training, a mask is used to select confident pseudo-labels. Specifically, a threshold *τ* is used to identify high-confidence predictions from weakly augmented data and generate masks as follows:

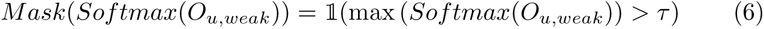

For each *i*th sample, pseudo label *P* ^(*i*)^ is generated by selecting the class with the highest logit from 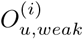:

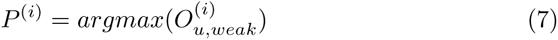

Finally, the unsupervised loss (i.e., consistency loss) *ℒ_u_* is computed as follows:

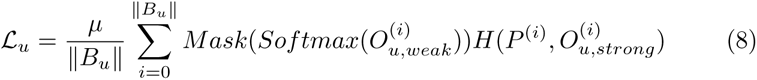

Here, *µ* represents the weight assigned to the unsupervised loss and *H*(*P, O_u,strong_*) denotes the cross-entropy loss between *P* and the logits of the strongly augmented samples.

### Data augmentations

We observed that existing data augmentations [11, 14] are insufficient for preventing overfitting of the model and achieving high performance in RUN-DVC (see Supplementary Fig. S3). Therefore, we propose a set of new augmentations and augmentation policies that are tailored for DNA sequencing datasets.

#### List of data augmentations

In the following, we provide overviews of the augmentations we proposed. Visualized examples of our augmentations are provided in Supplementary Fig. S5.

- **Subsampling:** Sub-sampling augmentation drops a specified portion of reads in the 3-dimensional tensor, with the proportion of reads to be removed specified as an input and the selection of reads to be dropped made randomly using a uniform distribution. This helps the DNN model become more robust against variation in the number of reads aligned in the candidate variant region.
- **Shifting and flipping:** Vertical shifting proved effective in preventing overfitting and improving the overall performance of RUN-DVC, while horizontal shifting did not lead to a performance increase. We chose not to use vertical flipping, as it reverses the order of aligned reads, which the model does not encounter at inference. Similarly, horizontal flipping was deemed unsuitable because it has the potential to alter variant labels [12, 40]. For example, in DNA sequencing, variants are often left-aligned, whereas most DNA sequencing processing tools output left-aligned variants. Thus, horizontal flipping would require changing left-aligned variants to right-aligned before flipping, which could alter the variant label.
- **Feature distortions:** Distortions were applied to different channels of the tensor, including reference bases, variants in the reads, base quality, mapping quality, and haplotype information. We randomly selected bases not located at the candidate variant’s position using a uniform distribution to distort the reference base channel, with the number of reference bases to change specified as input. We also introduced random false variants into the tensor using a uniform distribution to improve the model’s robustness to changes in the distribution of false variants. The number of false variants is given as input, and their positions are selected using a uniform distribution. We distorted mapping and base quality by setting mapping quality values to the maximum value or adding noise sampled from a normal distribution to base quality values. For haplotype information, we dropped the information by setting all values in the channel to the value used for the unphased case.

#### Weak and strong data augmentations

Our study incorporated two distinct data augmentation strategies within RUN-DVC, as delineated in Supplementary Algorithm. S1. The first strategy, termed weak augmentation, implemented subsampling and vertical shifting with a probability of 50%. In contrast, the second strategy, referred to as strong augmentation, made use of the random augmentation (RandAugment) policy [41], allowing for the selection of augmentations from feature distortions alongside subsampling and vertical shifting augmentations. The number of augmentations selected from feature distortions was capped at two, a limit we observed as necessary to achieve higher performance and circumvent overfitting during the training process.

### Training and hyperparameters

The stochastic weight averaging [42, 43] technique is used in the training. A running average of model weights is kept during training that is used for validation and model saving. Specifically, the average of model weights is updated every 32 iterations with a decay ratio of 0.99.

In all experiments, the same hyperparameters are used unless it is specified (e.g., for evaluation in ONT datasets). Radam [44] optimizer is used with initial learning rate *l_r_* = 3*e^−^*^5^. Additionally, an exponential learning rate scheduler was employed, reducing the learning rate by a factor of 0.97 per epoch. The number of training epochs is set to 50 epochs where the number of iterations in one epoch is determined by the number of batches in the source dataset. We used a batch size of 1000 for the labeled and unlabeled datasets, 0.9 for the confidence threshold in masking, 0.05 for the weight of unsupervised loss, and 1*e^−^*^5^ for the weight decay. For all evaluations, we train models from scratch to focus on evaluating the performance of UDA and SSDA themselves. The best model is selected based on the validation loss in the source domain.

## Code availability

RUN-DVC is open-source software (BSD 3-Clause license), available at https://github.com/kaist-ina/RUN-DVC.

## Data availability

The links to the reference genomes, truth variants, benchmarking materials, and sequencing datasets, and commands used for DeepVariant and Clair3, are available in the Supplementary Materials.

## References

[1] Li, H. Toward better understanding of artifacts in variant calling from high-coverage samples. Bioinformatics 30 (20), 2843–2851 (2014) .

[2] Carvalho, C. & Lupski, J. R. Mechanisms underlying structural variant formation in genomic disorders. Nature Reviews Genetics 17 (4), 224–238 (2016) .

[3] Yi, K. & Ju, Y. S. Patterns and mechanisms of structural variations in human cancer. Experimental & molecular medicine 50 (8), 1–11 (2018) .

[4] Metzker, M. L. Sequencing technologies—the next generation. Nature reviews genetics 11 (1), 31–46 (2010) .

[5] Head, S. R. et al. Library construction for next-generation sequencing: overviews and challenges. Biotechniques 56 (2), 61–77 (2014) .

[6] Amarasinghe, S. L. et al. Opportunities and challenges in long-read sequencing data analysis. Genome biology 21 (1), 1–16 (2020) .

[7] Cortés-Ciriano, I., Gulhan, D. C., Lee, J. J.-K., Melloni, G. E. & Park, P. J. Computational analysis of cancer genome sequencing data. Nature Reviews Genetics 23 (5), 298–314 (2022) .

[8] Zook, J. M. et al. Integrating human sequence data sets provides a resource of benchmark snp and indel genotype calls. Nature biotechnology 32 (3), 246–251 (2014) .

[9] Zook, J. M. et al. An open resource for accurately benchmarking small variant and reference calls. Nature biotechnology 37 (5), 561–566 (2019) .

[10] Baid, G., et al. An extensive sequence dataset of gold-standard samples for benchmarking and development. bioRxiv (2020).

[11] Poplin, R. et al. A universal snp and small-indel variant caller using deep neural networks. Nature biotechnology 36 (10), 983–987 (2018) .

[12] Krusche, P. et al. Best practices for benchmarking germline small-variant calls in human genomes. Nature biotechnology 37 (5), 555–560 (2019) .

[13] Shafin, K. et al. Haplotype-aware variant calling with pepper-margin-deepvariant enables high accuracy in nanopore long-reads. Nature methods 18 (11), 1322–1332 (2021) .

[14] Zheng, Z. et al. Symphonizing pileup and full-alignment for deep learning-based long-read variant calling. Nature Computational Science 2 (12), 797–803 (2022) .

[15] Trost, B. et al. Impact of dna source on genetic variant detection from human whole-genome sequencing data. Journal of medical genetics 56 (12), 809–817 (2019) .

[16] Dong, Z. et al. Development of coupling controlled polymerizations by adapter-ligation in mate-pair sequencing for detection of various genomic variants in one single assay. DNA Research 26 (4), 313–325 (2019) .

[17] Ahsan, M. U., Liu, Q., Fang, L. & Wang, K. Nanocaller for accurate detection of snps and indels in difficult-to-map regions from long-read sequencing by haplotype-aware deep neural networks. Genome biology 22 (1), 1–33 (2021) .

[18] Meng, J., Victor, B., He, Z., Liu, H. & Jiang, T. Deepssv: detecting somatic small variants in paired tumor and normal sequencing data with convolutional neural network. Briefings in Bioinformatics 22 (4), bbaa272 (2021) .

[19] Khazeeva, G. et al. Denovocnn: a deep learning approach to de novo variant calling in next generation sequencing data. Nucleic acids research 50 (17), e97–e97 (2022) .

[20] Krishnamachari, K. et al. Accurate somatic variant detection using weakly supervised deep learning. Nature Communications 13 (1), 4248 (2022) .

[21] Chen, N.-C. et al. Improving variant calling using population data and deep learning. BMC bioinformatics 24 (1), 1–15 (2023) .

[22] Sahraeian, S. M. E. et al. Deep convolutional neural networks for accurate somatic mutation detection. Nature communications 10 (1), 1–10 (2019) .

[23] Luo, R., Sedlazeck, F. J., Lam, T.-W. & Schatz, M. C. A multi-task convolutional deep neural network for variant calling in single molecule sequencing. Nature communications 10 (1), 1–11 (2019) .

[24] Luo, R. et al. Exploring the limit of using a deep neural network on pileup data for germline variant calling. Nature Machine Intelligence 2 (4), 220–227 (2020) .

[25] Kolesnikov, A. et al. Deeptrio: variant calling in families using deep learning. bioRxiv (2021) .

[26] Zook, J. M. et al. Extensive sequencing of seven human genomes to characterize benchmark reference materials. Scientific data 3 (1), 1–26 (2016) .

[27] Zook, J. M. et al. A robust benchmark for detection of germline large deletions and insertions. Nature biotechnology 38 (11), 1347–1355 (2020) .

[28] Wagner, J. et al. Benchmarking challenging small variants with linked and long reads. Cell Genomics 2 (5), 100128 (2022). https://doi.org/https://doi.org/10.1016/j.xgen.2022.100128.

[29] Wang, T. et al. The human pangenome project: a global resource to map genomic diversity. Nature 604 (7906), 437–446 (2022) .

[30] Consortium, . G. P. et al. A global reference for human genetic variation. Nature 526 (7571), 68 (2015) .

[31] Ball, M. P. et al. Harvard personal genome project: lessons from participatory public research. Genome medicine 6 (2), 1–7 (2014) .

[32] Berthelot, D., Roelofs, R., Sohn, K., Carlini, N. & Kurakin, A. Adamatch: A unified approach to semi-supervised learning and domain adaptation. arXiv preprint arXiv:2106.04732 (2021) .

[33] Saito, K., Kim, D., Sclaroff, S., Darrell, T. & Saenko, K. Semi-supervised domain adaptation via minimax entropy. Proceedings of the IEEE/CVF International Conference on Computer Vision 8050–8058 (2019) .

[34] Saito, K., Kim, D., Sclaroff, S. & Saenko, K. Universal domain adaptation through self supervision. Advances in neural information processing systems 33, 16282–16292 (2020) .

[35] Hommelsheim, C. M., Frantzeskakis, L., Huang, M. & Ülker, B. Pcr amplification of repetitive dna: a limitation to genome editing technologies and many other applications. Scientific reports 4 (1), 5052 (2014) .

[36] Vilov, S. & Heinig, M. Deepsom: a cnn-based approach to somatic variant calling in wgs samples without a matched normal. Bioinformatics 39 (1), btac828 (2023) .

[37] Nvidia parabricks retraining tool (2023). https://catalog.ngc.nvidia.com/orgs/nvidia/collections/claraparabricks/entities.

[38] Sohn, K. et al. Fixmatch: Simplifying semi-supervised learning with consistency and confidence. Advances in neural information processing systems 33, 596–608 (2020) .

[39] Xie, Q., Dai, Z., Hovy, E., Luong, T. & Le, Q. Unsupervised data augmentation for consistency training. Advances in Neural Information Processing Systems 33, 6256–6268 (2020) .

[40] Tan, A., Abecasis, G. R. & Kang, H. M. Unified representation of genetic variants. Bioinformatics 31 (13), 2202–2204 (2015) .

[41] Cubuk, E. D., Zoph, B., Shlens, J. & Le, Q. V. Randaugment: Practical automated data augmentation with a reduced search space. Proceedings of the IEEE/CVF conference on computer vision and pattern recognition workshops 702–703 (2020) .

[42] Tarvainen, A. & Valpola, H. Mean teachers are better role models: Weight-averaged consistency targets improve semi-supervised deep learning results. Advances in neural information processing systems 30 (2017) .

[43] Izmailov, P., Podoprikhin, D., Garipov, T., Vetrov, D. & Wilson, A. G. Averaging weights leads to wider optima and better generalization. arXiv preprint arXiv:1803.05407 (2018) .

[44] Liu, L. et al. On the variance of the adaptive learning rate and beyond. arXiv preprint arXiv:1908.03265 (2019) .

