## Supplementary Material for "Generalizing deep variant callers via domain adaptation and semi-supervised learning"

### CONTENTS

**1 Details of inputs for CNN and RNN model.** **2**

**2 Commands** **3**

**3 Links for datasets** **3**

**4 Details of training datasets of Clair3 and DeepVariant (PEPPER)** **3**

### LIST OF ALGORITHMS

S1 Data Augmentation Procedure . . . . . 4
S2 RUN-DVC Loss Function . . . . . 4

### 1. DETAILS OF INPUTS FOR CNN AND RNN MODEL.

**Input tensor for CNN model.** The 3-dimensional tensor comprises multiple channels, with each channel providing information about the candidate variant region. Although much of our tensor design is comparable to that used in Clair3, we made three critical modifications. First, we assign the mapping quality value not just to positions with aligned reads, but also to positions where a deletion is indicated. This modification is intended to circumvent a potential confounding scenario where the mapping information is zero when the deletion spans beyond the window size of the tensor. Second, in the target variant channel, we allocate the allele frequency value solely to the position of the candidate variant. This adjustment was made to enable the horizontal shift data augmentation techniques. Third, we changed the values used for encoding bases and strand information. Our refined three-dimensional tensor comprises eight distinct channels. These channels encode information about reference bases, observed variants in reads, strand information, mapping quality, base quality, the position of candidate variants, the bases involved in insertions, and phasing information. However, in the case of short-read sequencing data, the phasing channel is omitted, resulting in a total of seven channels. The input size for our tensor differs depending on the sequencing platform. For short-read sequencing platforms and the PacBio sequencing platform, we utilized an input tensor with dimensions of 33 columns by 55 rows. However, for the ONT sequencing platform, we employed an input tensor of 33 columns by 89 rows to account for the increased read depth. By default, we assign a value of zero to all positions where either a deletion is indicated or no reads are mapped.

- **Reference bases:** Each position of the aligned read is assigned an integer based on the reference base at that position (A: 75, C: -50, G: 50, T: -75).
- **Mutations:** Positions differing from the reference base are assigned an integer based on the type of variant present. SNPs are assigned an integer based on the alternative base at the position using the same base-value mapping as in the reference bases channel. INDEL variants are assigned -25 and 25, respectively, at positions corresponding to the starting location of the INDEL on the left end.
- **Strand information:** Each read position is assigned an integer depending on the strand the read is aligned to: 50 for the forward strand and -50 for the reverse strand.
- **Mapping quality:** An integer between 0 and 100 is assigned to all mapped read positions, scaled up from the original Phred mapping score (range 0 to 60), and capped at 60 if it exceeds that value.
- **Base quality:** Each aligned base is assigned an integer between 0 and 100, based on the scaled-up Phred base quality score (original range 0 to 40) and capped at 40 if it exceeds that value.
- **Target variant:** The candidate variant position (default is the column center) is assigned an integer between 0 and 100, indicating the percentage of reads supporting the specific mismatch pattern. For instance, if 10 out of 20 reads support the reference allele "A", 5 support the alternative "C", 3 support "T", and 2 support an insertion "AT", the integer assigned would be 0, 25, 15, and 10, respectively.
- **Insertion bases:** Inserted bases are encoded following each insertion's starting position, using the same base-value mapping as the reference bases channel.
- **Phasing information:** All positions of each read are assigned an integer value based on phasing: -50 for HP1, 20 for unphased, and 50 for HP2 reads. If reads are phased, they are sorted in the order "unphased, HP1, HP2" across all eight channels.

**Input tensor for RNN model.** The pileup input tensor consists of 594 integers, representing 33 genome positions with 18 features at each position. These features include counts of read support for the four nucleotides (A, C, G, T), insertions ($I_1$ and $I$), deletions ($D_1$ and $D$), and $R$ on both the positive strand (+) and negative strand (-). The presence of a '1' superscript indicates that only the indel with the highest read support is counted if there are multiple indels at a given candidate site (i.e., all indels are counted if there is no '1' superscript). The 'R' indicates the following positions of an indel.

### 2. COMMANDS

Please see the GitHub for commands used for RUN-DVC. We provide scripts regarding dataset generation, training, and variant calling which heavily make use of GNU Parallel [1].

**Command used for DeepVariant**

```
docker run google/deepvariant:1.5.0 /opt/deepvariant/bin/run_deepvariant -model_type
WGS -ref {Reference} -reads {BAM} -output_vcf {OutputFile} -num_shards {ThreadNum}
-regions {BED}
```

**Command used for PEPPER**

```
sudo docker run  -u ‘id -u $USER‘:‘id -g $USER‘  -ipc=host  kishwars/pepper_deepvariant:r0.8
run_pepper_margin_deepvariant call_variant  -b "${BAM_FILE}"  -f "/data/Homo_sapiens_assembly38.fasta"
-o "/output/"  -regions /data/HG003_GRCh38_1_22_v4.2.1_benchmark.bed  -p "${DATASET}"
-t "${THREADS}  -ont_r9_guppy5_sup
```

**Command used for VCF report in DeepVariant**

```
docker run google/deepvariant:1.5.0 /opt/deepvariant/bin/vcf_stats_report -input_vcf
${INPUT_VCF} -outfile_base ${OUTPUT}
```

**Command used for Clair3**

```
docker run hkubal/clair3:latest /opt/bin/run_clair3.sh -model_path="/opt/models/{MODEL_NAME}"
-ref_fn={Reference} -bam_fn={BAM} -output={OutputFile} -threads={ThreadNum} -bed_fn={BED}
-platform={platform}
```

### 3. LINKS FOR DATASETS

**Genome stratification file:**
https://ftp-trace.ncbi.nlm.nih.gov/giab/ftp/release/genome-stratifications/

**GIAB truth variant sets:**
https://ftp-trace.ncbi.nlm.nih.gov/giab/ftp/release/

**GIAB sequencing datasets:**
https://ftp-trace.ncbi.nlm.nih.gov/giab/ftp/data/

**Google sequencing datasets:**
https://console.cloud.google.com/storage/browser/brain-genomics-public

**Human Pangenome Reference Consortium datasets:**
https://s3-us-west-2.amazonaws.com/human-pangenomics/index.html?prefix=NHGRI_UCSC_panel/

### 4. DETAILS OF TRAINING DATASETS OF CLAIR3 AND DEEPVARIANT (PEPPER)

The training datasets used for Clair3 and DeepVariant can be found in below links.
DeepVariant for PacBio dataset: https://github.com/google/deepvariant/blob/r1.5/docs/deepvariant-details-training-data.md
Clair3 for PacBio dataset: https://github.com/HKU-BAL/Clair3/blob/main/docs/training_data.md
Clair3 for ONT dataset: https://github.com/HKU-BAL/Clair3/blob/main/docs/guppy5_20220113.md

**Algorithm S1.** Data Augmentation Procedure

```
data ← Sequencing data in 3D tensor
List_Aug ← ["row_drop", "vertical_shift"]
List_Feature_Aug ← ["Reference", "Mapping_quality", "Base_quality", "Target_variants",
"Phasing"]
if weak augmentation then
    if random.random() < 0.5 then
        data.ApplyAug(Intensity=0.3, List_Aug)
else if strong augmentation then
    data.ApplyAug(Intensity=0.7, List_Aug)
    data.ApplyRandaug(Intensity=0.5, Num_Aug=2, List_Feature_Aug)
```

**Algorithm S2.** RUN-DVC Loss Function

```
procedure RUNDVC_LOSS(label_s, lc, lspp, source_total)
    /*
    param label_s: labels of the source domain batch
    param lc: logits obtained from the combined batch (source + target)
    param lspp: logits obtained from the source batch (only source)
    param source_total: size of the weakly and strongly augmented source batch
    */
    source_batch_size ← source_total//2
    target_batch_size ← (lc.size(0) − source_total)//2
    lsp ← lc[: source_total]
    if USE_RLI then
        lambd ← torch.rand_like(lsp)
        final_logits_source ← (lambd ∗ lsp) + (1 − lambd) ∗ lspp
    else
        final_logits_source ← lsp
    source_loss ← CrossEntropyLoss(final_logits_source, label_s)
    if USE_SSL then
        logits_target_weak ← lc[source_total : −target_batch_size]
        logits_target_strong ← lc[−target_batch_size :])
        mask ← Softmax(logits_target_weak) > Confidence_Value
        pseudo_labels ← argmax(logits_target_weak)
        target_loss ← CrossEntropy(logits_target_strong, pseudo_labels)
        target_loss ← mask × target_loss
        Return source_loss, target_loss
    else
        Return source_loss
```

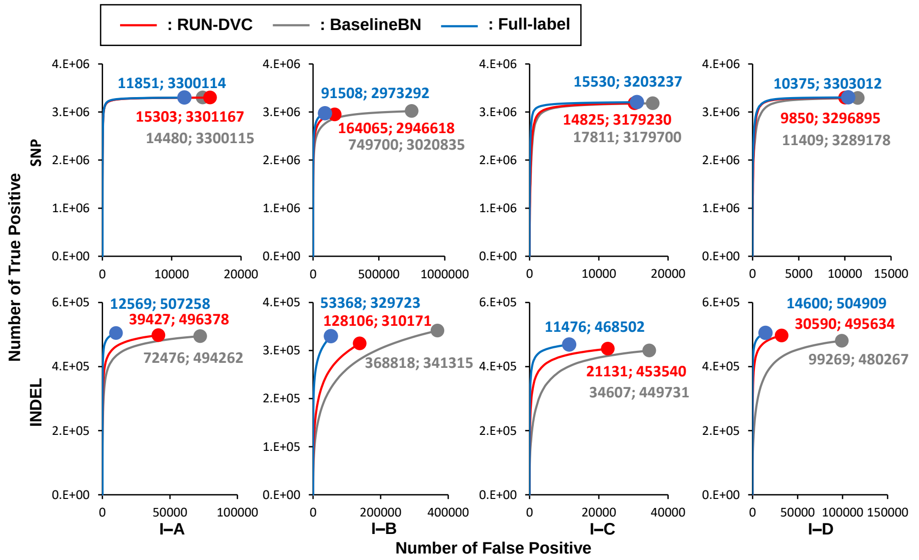

(a) A number of true positives and false positives. The number of false positives and true positives for all PASS calls is marked with a solid circle and a label (the number of false positives; the number of true positives).

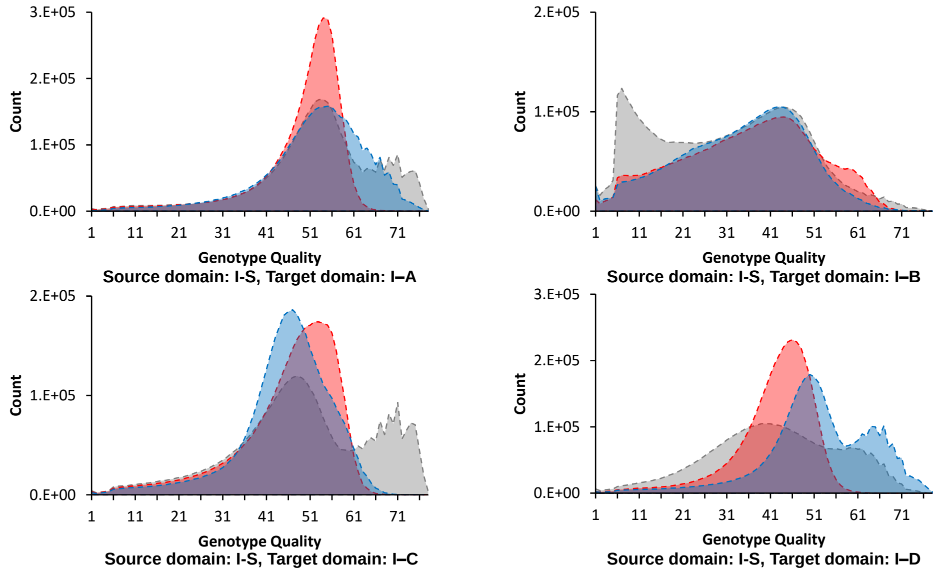

(b) Genotype quality distribution results.

**Fig. S1.** Performance analysis of RUN-DVC, *BaselineBN*, and Full-label on I-A, I-B, I-C, and I-D datasets in UDA setting.

| Name | Sequencing Platform | Sample and Library preparation method |
| --- | --- | --- |
| I-Source | Illumina Novaseq PCR-free | [2] |
| I-A | Illumina Novaseq PCR-plus | [2] |
| I-B | Illumina Hiseq2500 | https://ftp-trace.ncbi.nlm.nih.gov/giab/ftp/data/AshkenazimTrio/HG002_NA24385_son/10XGenomics/README_10Xgenomes.txt |
| I-C | BGISEQ-500 | https://ftp-trace.ncbi.nlm.nih.gov/giab/ftp/data/NA12878/stLFR/readme.txt |
| I-D | Illumina HiseqX PCR-plus | [2] |
| P-Source | PacBio CCS Sequel II | https://ftp-trace.ncbi.nlm.nih.gov/giab/ftp/data/AshkenazimTrio/HG002_NA24385_son/PacBio_CCS_15kb_20kb_chemistry2/GRCh38/README.txt<br>https://ftp-trace.ncbi.nlm.nih.gov/giab/ftp/data/AshkenazimTrio/HG002_NA24385_son/PacBio_SequelII_CCS_11kb/HG002_GRCh38/README.txt |
| P-A | PacBio CCS Sequel I | https://ftp-trace.ncbi.nlm.nih.gov/giab/ftp/data/AshkenazimTrio/HG002_NA24385_son/PacBio_CCS_10kb/GRCh38_no_alt_analysis/README.txt<br>https://ftp-trace.ncbi.nlm.nih.gov/giab/ftp/data/AshkenazimTrio/HG002_NA24385_son/PacBio_CCS_15kb/GRCh38_no_alt_analysis/README.txt |
| O-Source, O-A | ONT datsets | http://www.bio8.cs.hku.hk/guppy5_data/<br>https://github.com/HKU-BAL/Clair3/blob/main/docs/guppy5_20220113.md#how-to-use-the-guppy5-model |

**Table S1.** Details of sequencing datasets used.

| | | I-A | | | I-B | | |
|---|---|---|---|---|---|---|---|
| Method | Variant type | Recall | Precision | F1-score | Recall | Precision | F1-score |
| BaselineBN | Indel | 94.06% | 87.21% | 90.51% | 66.22% | 48.06% | 55.70% |
| | SNP | 99.14% | 99.56% | 99.35% | 90.76% | 80.12% | 85.11% |
| Clair3 | Indel | 94.53% | 87.18% | 90.71% | 67.93% | 41.68% | 51.66% |
| | SNP | 99.18% | 99.52% | 99.35% | 91.75% | 57.94% | 71.03% |
| DeepVariant | Indel | 96.83% | 96.56% | 96.69% | 66.91% | 55.92% | 60.92% |
| | SNP | 99.20% | 99.84% | 99.52% | 90.85% | 94.21% | 92.50% |
| RUN-DVC (UDA) | Indel | 94.65% | 92.28% | 93.45% | 61.22% | 69.42% | 65.06% |
| | SNP | 99.17% | 99.53% | 99.35% | 88.54% | 94.69% | 91.51% |
| Full-label | Indel | 96.70% | 97.58% | 97.14% | 64.62% | 86.07% | 73.82% |
| | SNP | 99.14% | 99.64% | 99.39% | 89.33% | 97.01% | 93.02% |

**Table S2.** Variant calling result (PASS calls) in HG002 sample of I-A and I-B.

### REFERENCES

1. O. Tange *et al.*, "Gnu parallel-the command-line power tool," The USENIX Mag. **36**, 42–47 (2011).
2. G. Baid, M. Nattestad, A. Kolesnikov, S. Goel, H. Yang, P.-C. Chang, and A. Carroll, "An extensive sequence dataset of gold-standard samples for benchmarking and development," bioRxiv (2020).

| | | I-C | | | I-D | | |
|---|---|---|---|---|---|---|---|
| Method | Variant type | Recall | Precision | F1-score | Recall | Precision | F1-score |
| BaselineBN | Indel | 86.29% | 92.85% | 89.45% | 91.42% | 82.87% | 86.93% |
| | SNP | 95.50% | 99.44% | 97.43% | 98.81% | 99.65% | 99.23% |
| Clair3 | Indel | 88.63% | 92.93% | 90.73% | 94.05% | 85.94% | 89.81% |
| | SNP | 95.93% | 99.38% | 97.62% | 99.32% | 99.40% | 99.36% |
| DeepVariant | Indel | 86.96% | 95.91% | 91.22% | 96.81% | 97.58% | 97.19% |
| | SNP | 89.20% | 99.67% | 94.15% | 99.27% | 99.81% | 99.54% |
| RUN-DVC (UDA) | Indel | 87.35% | 95.27% | 91.14% | 94.71% | 93.85% | 94.28% |
| | SNP | 95.49% | 99.52% | 97.47% | 99.04% | 99.70% | 99.37% |
| Full-label | Indel | 89.88% | 97.61% | 93.59% | 96.37% | 97.19% | 96.78% |
| | SNP | 96.20% | 99.52% | 97.83% | 99.22% | 99.69% | 99.45% |

**Table S3.** Variant calling result (PASS calls) in HG002 sample of I-C and I-D.

| | | P-A | | | O-A | | |
|---|---|---|---|---|---|---|---|
| Method | Variant type | Recall | Precision | F1-score | Recall | Precision | F1-score |
| BaselineBN | Indel | 97.14% | 96.22% | 96.68% | 74.01% | 80.42% | 77.08% |
| | SNP | 99.39% | 99.93% | 99.66% | 99.61% | 99.59% | 99.60% |
| Clair3 | Indel | 97.29% | 95.34% | 96.31% | 71.32% | 85.46% | 77.75% |
| | SNP | 99.88% | 99.92% | 99.90% | 99.62% | 99.66% | 99.64% |
| DeepVariant | Indel | 97.31% | 97.21% | 97.26% | 73.40% | 83.10% | 77.95% |
| | SNP | 99.90% | 99.95% | 99.93% | 99.66% | 99.76% | 99.71% |
| RUN-DVC (UDA) | Indel | 97.44% | 97.03% | 97.23% | 73.87% | 81.21% | 77.37% |
| | SNP | 99.42% | 99.93% | 99.68% | 99.60% | 99.58% | 99.59% |
| Full-label | Indel | 98.48% | 98.69% | 98.58% | 76.40% | 83.72% | 79.89% |
| | SNP | 99.56% | 99.93% | 99.75% | 99.63% | 99.58% | 99.60% |

**Table S4.** Variant calling result (PASS calls) in HG002 sample of P-A and O-A.

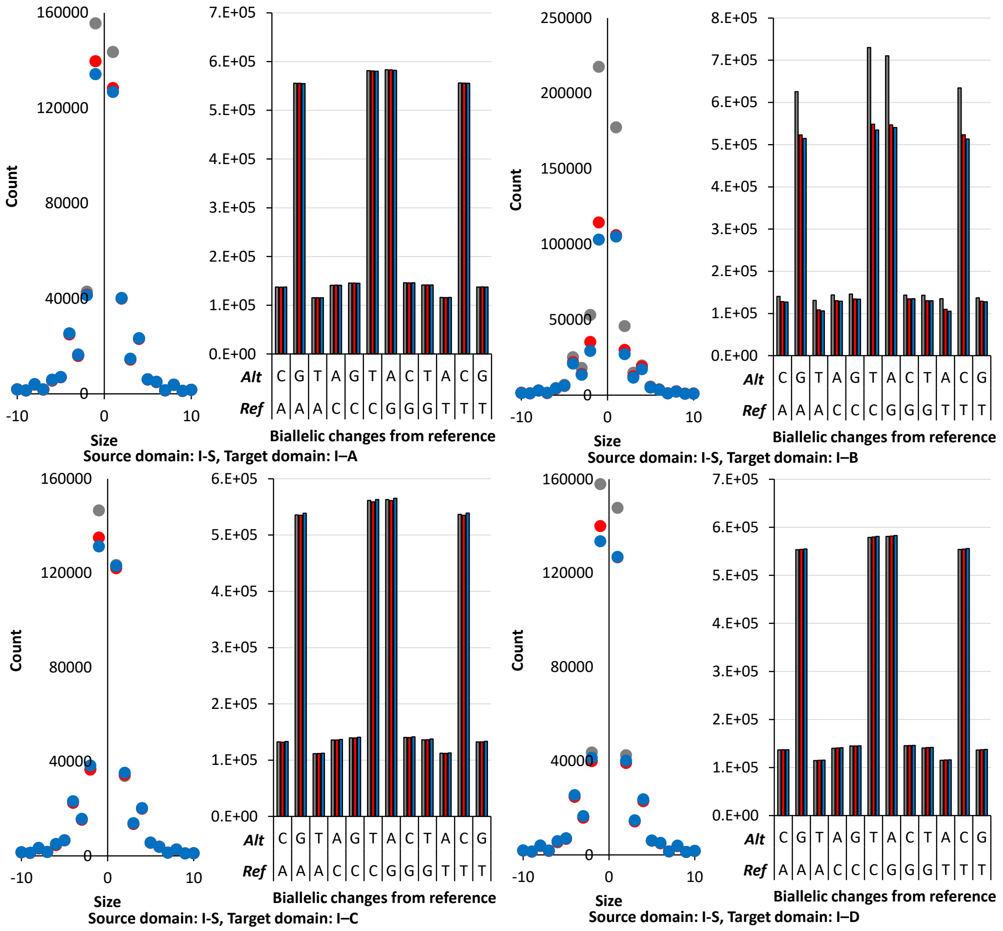

**Fig. S2.** Analysis on disagreements between RUN-DVC, *BaselineBN*, and *Full-label* in UDA setting.

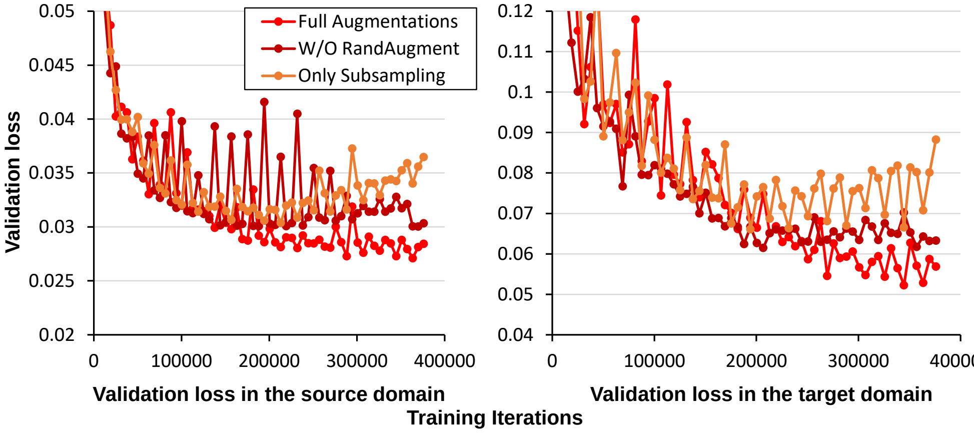

**Fig. S3.** An ablation study demonstrating the impact of varying data augmentation strategies on RUN-DVC in a UDA setting (Source: I-Source, Target: I-A).
The "Only Subsampling" method applies subsampling as the sole augmentation for both weak and strong augmentation policies. "W/O RandAugment" incorporates both subsampling and vertical shifting in weak and strong augmentation policies, excluding RandAugment. "Full Augmentations" extends the "W/O RandAugment" by additionally including RandAugment in the strong augmentation policy.

| Dataset | Model type | ($\gamma = 0$) | ($\gamma = 0.5$) | ($\gamma = 1$) |
|---|---|---|---|---|
| I-A | RNN | 99.80% | 99.91% | 99.91% |
| I-B | RNN | 92.40% | 97.61% | 98.90% |
| | CNN | 94.02% | 97.69% | 98.88% |
| I-C | RNN | 99.64% | 99.90% | 99.90% |
| P-A | RNN | 99.91% | 99.92% | 99.92% |

**Table S5.** Accuracy analysis of binary classification for true variant and non-variant using RNN and CNN model.

| Dataset | Variant type | Recall | Precision | F1-score |
|---|---|---|---|---|
| I-A | Indel | 83.03% | 70.75% | 76.40% |
| | SNP | 99.05% | 99.12% | 99.08% |
| I-B | Indel | 57.94% | 49.48% | 53.38% |
| | SNP | 87.04% | 87.08% | 87.06% |
| I-C | Indel | 40.80% | 87.20% | 55.59% |
| | SNP | 34.16% | 95.50% | 50.33% |
| P-A | Indel | 83.97% | 90.42% | 87.08% |
| | SNP | 99.58% | 99.63% | 99.60% |

**Table S6.** Accuracy of variant calling using RNN model.

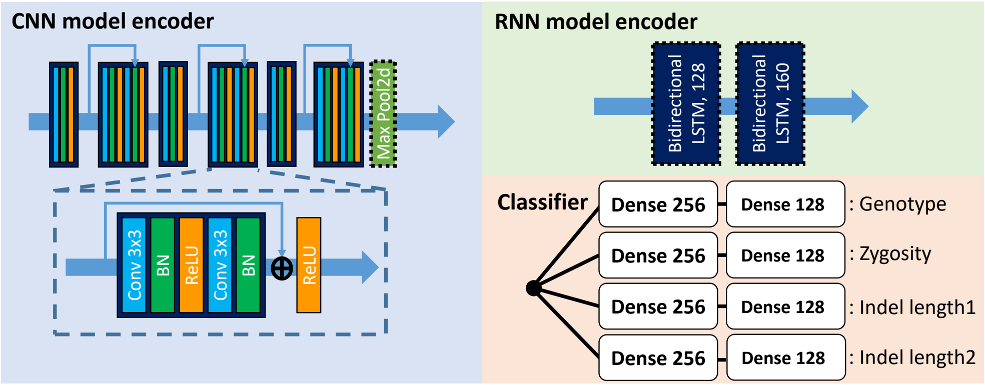

**Fig. S4.** Model architecture of the encoder and classifier.

**Table S7.** Performance of RUN-DVC and Baseline methods under various quantities of labeled datasets.

| Data | Metric | Method | Number of Labeled Datasets (million) | | | |
|---|---|---|---|---|---|---|
| | | | 0.4 | 0.8 | 1.6 | 3.2 |
| I-A | Validation Loss | RUNDVC | 0.03769 | 0.035506 | 0.033981 | 0.032647 |
| | Validation Loss | BaselineBN | 0.044177 | 0.040084 | 0.036158 | 0.033772 |
| | F1-score (INDEL) | RUNDVC | 0.959551 | 0.965844 | 0.967458 | 0.970639 |
| | F1-score (INDEL) | Baseline | 0.953517 | 0.959907 | 0.964962 | 0.968631 |
| | F1-score (SNP) | RUNDVC | 0.993561 | 0.993741 | 0.993702 | 0.993811 |
| | F1-score (SNP) | Baseline | 0.993608 | 0.993665 | 0.993693 | 0.993779 |
| I-B | Validation Loss | RUNDVC | 0.112521 | 0.106701 | 0.10008 | 0.096151 |
| | Validation Loss | BaselineBN | 0.141616 | 0.118686 | 0.104717 | 0.098638 |
| | F1-score (INDEL) | RUNDVC | 0.693044 | 0.701516 | 0.712803 | 0.717854 |
| | F1-score (INDEL) | Baseline | 0.669771 | 0.694079 | 0.710827 | 0.717897 |
| | F1-score (SNP) | RUNDVC | 0.919777 | 0.920341 | 0.921952 | 0.923228 |
| | F1-score (SNP) | Baseline | 0.90728 | 0.914979 | 0.919946 | 0.921779 |
| I-A (seed2) | Validation Loss | RUNDVC | 0.037763 | 0.035852 | 0.034277 | 0.032966 |
| | Validation Loss | BaselineBN | 0.043844 | 0.039584 | 0.036641 | 0.033777 |
| | F1-score (INDEL) | RUNDVC | 0.960536 | 0.96486 | 0.968501 | 0.970342 |
| | F1-score (INDEL) | Baseline | 0.952172 | 0.959439 | 0.965232 | 0.96901 |
| | F1-score (SNP) | RUNDVC | 0.993628 | 0.993658 | 0.993729 | 0.993809 |
| | F1-score (SNP) | Baseline | 0.993585 | 0.993606 | 0.993663 | 0.993806 |
| I-B (seed2) | Validation Loss | RUNDVC | 0.110411 | 0.105141 | 0.100272 | 0.096536 |
| | Validation Loss | BaselineBN | 0.13249 | 0.118803 | 0.103799 | 0.099854 |
| | F1-score (INDEL) | RUNDVC | 0.69594 | 0.705068 | 0.711312 | 0.718372 |
| | F1-score (INDEL) | Baseline | 0.677013 | 0.694128 | 0.710965 | 0.71763 |
| | F1-score (SNP) | RUNDVC | 0.919744 | 0.921293 | 0.921882 | 0.922887 |
| | F1-score (SNP) | Baseline | 0.909666 | 0.91393 | 0.919964 | 0.921768 |

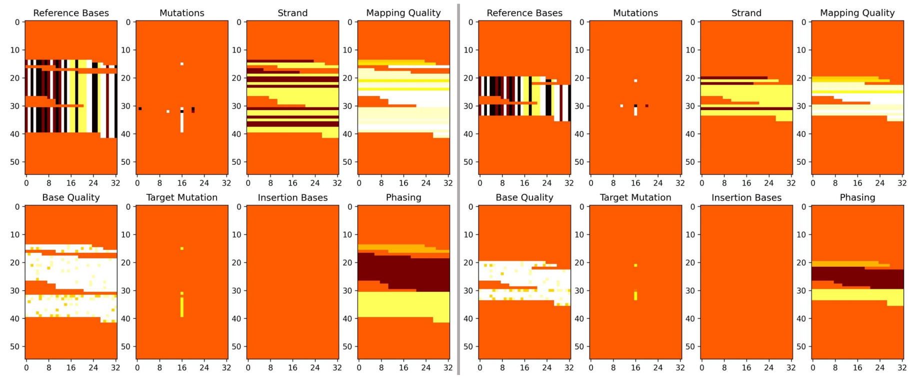

(a) Left: Original, Right: Down sampling

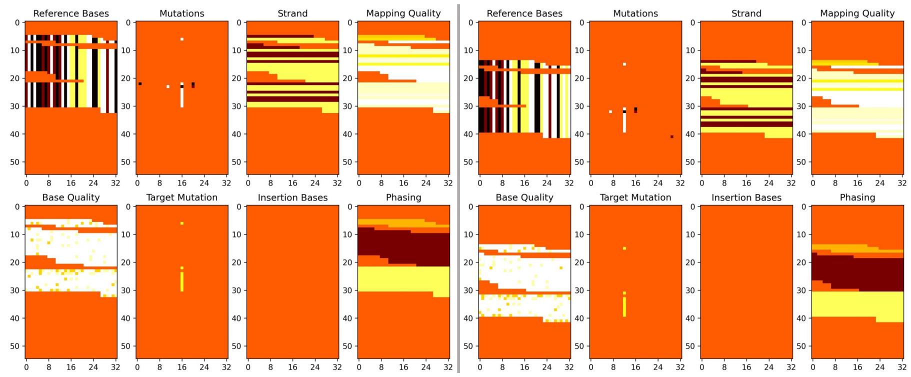

(b) Left: Vertical shift, Right: Horizontal shift

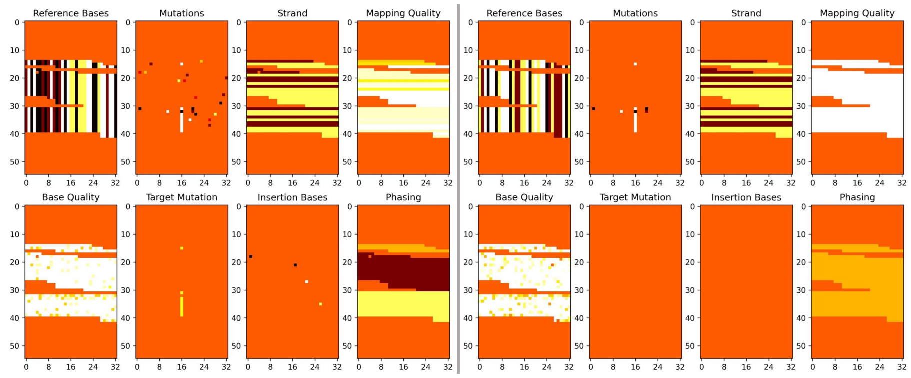

(c) Left: SNP/Indel variants, Right: Distortions to Reference bases, Mapping quality, Base quality, Target variant, and Phasing

**Fig. S5.** Overview of data augmentations
